# A multiplexed approach for genetic screening of human cells by electron microscopy uncovers a critical effector of mitochondrial cristae shape

**DOI:** 10.1101/2025.11.15.688601

**Authors:** Sarah Hassdenteufel, Yury Bykov, Benjamin Dubreuil, Katja Noll, Ofrah Faust, Roman Kamyshinsky, Maxim Itkin, Nir Cohen, Sergey Malitsky, Yoav Peleg, Shira Albeck, Karina von der Malsburg, Deborah Fass, Martin van der Laan, Maya Schuldiner

**Author notes:** Quantitative Cell Biology, Rhineland-Palatinate Technical University, Kaiserslautern 67663, Germany (Yury Bykov).

## Abstract

Mitochondria form complex and diverse membrane-architectures essential for their multiple functions. Whereas critical proteins sculpting mitochondrial membranes have been identified, the molecular basis for many key features remains enigmatic. Exploration of membrane ultrastructure, in general, is limited by the tradeoff between resolution and throughput: electron microscopy (EM) is essential to resolve their ultrastructure but lacks scalability for systematic functional discovery. To overcome this limitation, we developed a high-resolution screening pipeline for EM of multiplexed human cell pools, hMultiCLEM (<u>h</u>uman <u>Multi</u>plexed <u>C</u>orrelative <u>L</u>ight and <u>EM</u>). To showcase the power of hMultiCLEM we performed a genetic screen exploring mitochondrial ultrastructure. hMultiCLEM confirmed proposed cristae modulators and uncovered additional ones illuminating the protein networks driving cristae organization. Validation of candidates highlighted an intermembrane space (IMS) protein linked to Ménière’s disease, which we named MISHA (<u>M</u>itochondrial-IMS membrane-<u>SHA</u>pe-impacting protein). More broadly, hMultiCLEM transforms the EM field, enabling genetic/chemical screening in basic and medical research.

## Introduction

Mitochondria have a highly complex membrane architecture intertwined with their multitude of functions. The outer mitochondrial membrane (OMM) delimits the organelle, whereas the inner mitochondrial membrane (IMM) invaginates to form a unique ultrastructure, termed cristae, which harbor the machinery for ATP synthesis through oxidative phosphorylation (OXPHOS).

Cristae structures can be dynamically modulated by metabolic and developmental signals to accommodate cellular and organismal needs. At the cellular level, cristae abundance and morphology are actively controlled reflecting the energetic requirements and activity states of cells [1–3]. Heterogeneity in cristae morphology inside individual cells allows formation of subpopulations that support distinct mitochondrial functions [4]. At the organismal level, as cells differentiate during embryogenesis, mitochondria undergo specialization by changing their ultrastructure in a programmed manner. For example, mitochondria from brown adipocytes or cardiomyocytes have densely packed lamellar cristae to cater to high energy demands. Conversely, in hepatocytes, a reduced cristae area favors enzymatic processes taking place in the matrix [5].

Hence, it is no surprise that when mitochondria are impaired in defining and adjusting their ultrastructure optimal for a particular cell and organ, mitochondrial activity is impacted, and diseases can develop [6–9]. Mitochondria also respond to different disease states by altering their ultrastructure. Indeed, morphological changes have been documented in a multitude of neurodegenerative and cardiovascular diseases, though it is often not clear whether they are causal or consequential [10]. More generally, aging is accompanied by characteristic ultrastructural alterations in specific cell types [11]. Hence it is important to understand how mitochondria establish, and modulate [12,13], the diversity of their ultrastructure to deviate from the prototypic architecture.

A major step in understanding cristae organization was the discovery of the MICOS (<u>MI</u>tochondrial contact site and <u>C</u>ristae-<u>O</u>rganizing <u>S</u>ystem) complex [14–16]. The MICOS complex is localized at cristae junctions, where it stabilizes the sites of membrane invagination through oligomerization [17–19]. Two additional major components contribute to cristae shape: the F_1_Fo-ATP synthase complex residing at cristae tips and inducing high curvature through dimerization and oligomerization [20–23], and the dynamin-like GTPase OPA1, which remodels the shape and width of both tubular and lamellar invaginations through oligomerization of its short isoform [24,25]. All three well-characterized complexes share an intrinsic capacity to bend membranes and hence impact cristae organization directly [18,25–28].

An additional parameter is the lipid composition, which affects membrane curvature directly through differential shape of phospholipids and indirectly through stabilizing the protein complexes described above. Hence, proteins modulating the mitochondrial lipid composition affect cristae shape [29–33]. Physicochemical factors such as pH, ionic strength and mechanical forces further influence membrane packing and fluidity, so any protein impacting these parameters would contribute to the modulation of mitochondrial ultrastructure as well [13]. The regulators of these core proteins and lipids also impact mitochondrial ultrastructure through indirect effects. For example, proteases such as YME1L1 modulate the levels of OPA1 isoforms [34,35] with clear phenotypic alteration in cristae architecture. Together, the direct and indirect components create a dynamic network.

While these core components and regulators are well established [36–38], additional factors that affect cristae morphology are anticipated. Additional or alternative mechanisms for cristae formation must exist, since the absence of the three core players (MICOS, OPA1 and the ATP synthase) does not completely inhibit the invagination process [39]. Moreover, despite the ubiquity and evolutionary conservation of MICOS, cristae morphology varies extensively between species and across tissues [36,40], suggesting the presence of tissue-specific factors. Uncovering the full repertoire of cristae effectors will illuminate the networks and mechanisms that build and remodel mitochondrial ultrastructure in response to the changing requirements of a cell and an organism.

One major limitation that has slowed discovery of core modulators and regulators of this process is the requirement of imaging strategies that resolve ultrastructure such as electron microscopy (EM). While genetic screening at Light Microscopy (LM) resolution has uncovered many modulators of gross mitochondrial morphology [41], cristae are smaller than the diffraction limit of light and thus need EM to be resolved. However, EM is inherently low throughput due to the lengthy sample preparation and imaging procedures [42]. Hence, to date, it has not been possible to genetically screen for components impacting human mitochondrial ultrastructure, and their discovery has relied on non-systematic approaches. This limitation holds not only for mitochondrial ultrastructure but also for cellular ultrastructure in general and has hampered discovery in the cell biology of organelle membrane morphology.

To address this technological barrier, which broadly impacts organelle morphology studies, we created a new pipeline for screening human cells at EM resolution. This technology, hMultiCLEM (<u>H</u>uman <u>Multi</u>plexed <u>C</u>orrelative <u>L</u>ight and <u>EM</u>), uses fluorescent cell surface barcoding to multiplex samples, enabling visualization of dozens of samples across a few EM experiments. We showcase the utility of hMultiCLEM through a siRNA screen targeting poorly characterized mitochondrial proteins, to identify factors influencing mitochondrial ultrastructure. The hMultiCLEM screen identified characteristic phenotypes of the known core modulators as well as confirmed recently suggested new determinants [43–45]. Importantly, it revealed previously uncharacterized cristae effectors, among which secondary screens highlighted a soluble intermembrane space (IMS) protein which we named MISHA (<u>M</u>itochondrial-<u>I</u>MS membrane-<u>SHA</u>pe-impacting protein). MISHA is a newly-recognized protein of rapidly growing interest for which we found evidence of multiple structural states, including dimeric and higher-order assemblies. Our assays led us to propose a new mechanism by which MISHA influences cristae independently of core protein machineries and lipids, with possible implications for mitochondrial pathology, including familial Ménière’s disease, a polygenic inner-ear disorder with variable systemic manifestations [46].

More generally, the discovery of MISHA as a critical effector of mitochondrial ultrastructure, and the capacity to gain a systematic insight into a complex process such as cristae organization position hMultiCLEM as an effective approach for cellular exploration. hMultiCLEM thus advances EM imaging as a discovery tool for screening human samples using genetic or chemical approaches, and has potential applications in diagnostics, including the assessment of disease-associated mutations.

## Results

### Fluorescent conjugates enable surface barcoding for multiplexing of human cells

To enable screening of human cells at EM resolution, we extended our previously published approach for screening of *Saccharomyces cerevisiae* at EM resolution – MultiCLEM [47]. MultiCLEM exploits fluorescent barcoding of the yeast cell wall to enable pooling of multiple samples, thus reducing the labor of EM sample preparation. Genotype to phenotype decoding is then performed by the correlation of LM images (reporting on the sample barcode) and EM images (providing phenotypic information at ultrastructure resolution) (Fig. 1A). MultiCLEM in yeast relied on four-color barcoding using fluorophores conjugated to the lectin Concanavalin A (ConA), which binds the cell wall.

**Fig. 1:**
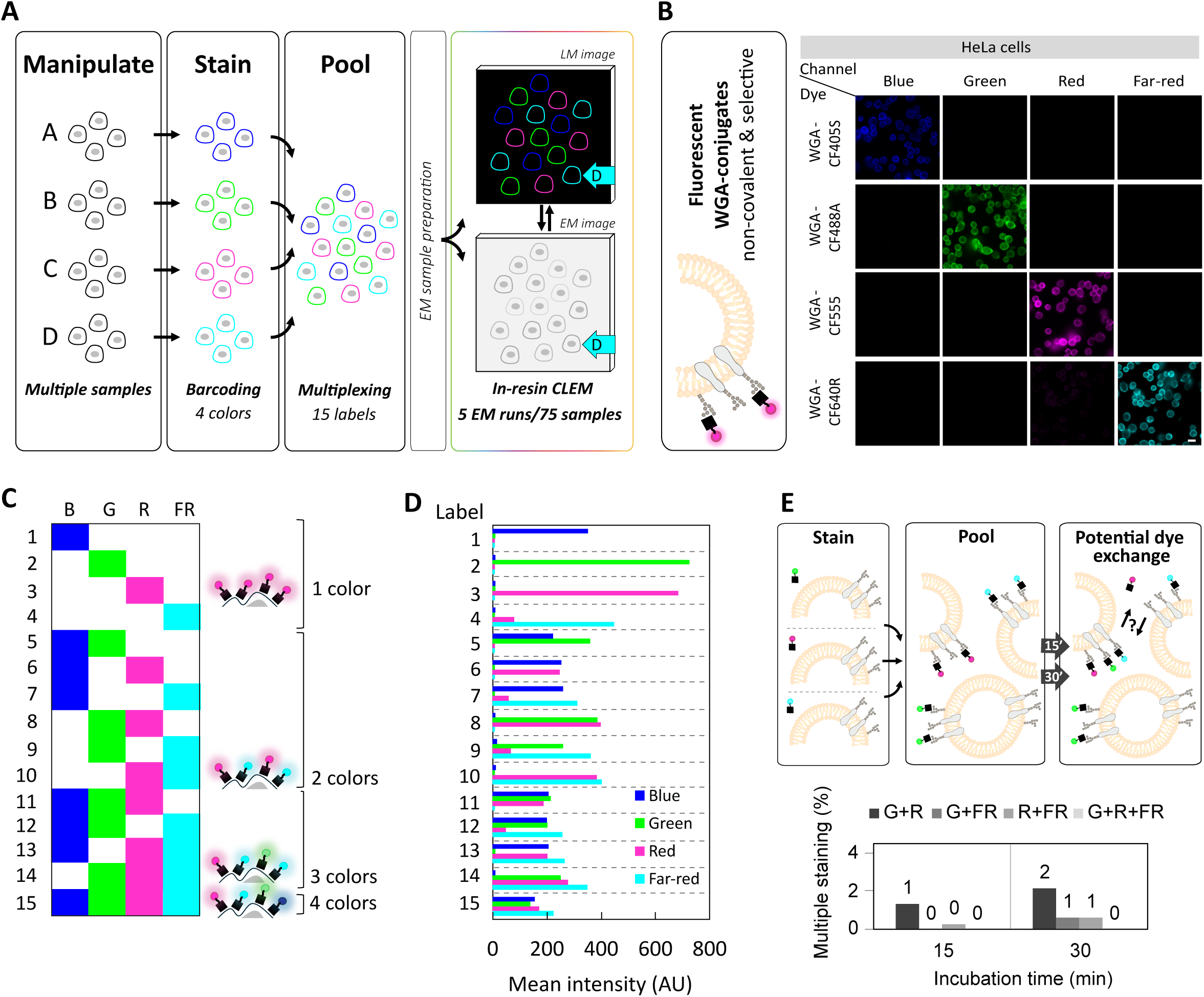
Fluorescent conjugates enable surface barcoding for multiplexing of human cells. **A)** Schematic of the barcoding and decoding process: cell populations are individually stained on the cell surface (barcoding) and pooled enabling multiplexing of 15 different samples. Following an in-resin Correlative Light and Electron Microscopy (CLEM) protocol for cell embedding and sectioning (EM sample preparation, time-limiting step), the pooled sample is imaged with light microscopy (LM) and electron microscopy (EM). Computational correlation of the two corresponding images enables barcode-decoding, i.e. assignment of cell identity to each cell. **B)** Fluorescent conjugates of Wheat Germ Agglutinin (WGA) enable barcoding: WGA non-covalently binds to selected carbohydrates on the plasma membrane. HeLa cells were detached and each sample individually stained with one of four WGA-conjugated fluorophores. Stained cell populations were automatically imaged in each of the four channels to assess signal separation (bleed-through). Scale bar: 20 μm. **C)** Combinatorial mixing of four different dyes achieves 15 labels (2^4^-1, zero color was omitted for reasons of accuracy, see Supplementary Fig. S1): Each cell of a respective label exposes one to four distinct emission colors (channels) on its surface: Blue (B), green (G), red (R) and far-red (FR). **D)** Dye concentrations were optimized to enable a robust four-channel signal profile: HeLa cells were detached and each sample individually stained with one of 15 combinations derived from four dyes, before automated imaging in the four channels. Following cell segmentation and quantification of fluorescence. Mean intensities are shown for each channel in arbitrary units (AU). **E)** Assessment of the misidentification potential due to dye exchange: HeLa cell samples were individually stained with one of three single colors, pooled, and incubated for prolonged times. At the indicated time points, a sample was automatically imaged and analyzed for dual or triple stained cells, shown in percentages.

While this method works well in yeast, ConA is not compatible with the glycan composition of the human plasma membrane, necessitating a different barcoding strategy. Moreover, human cells are larger and more heterogenous in shape and cannot be processed using the yeast-optimized MultiCLEM protocol. We therefore developed an optimal barcoding protocol for human cells [48] and created a new pipeline for correlation and identity-assignment of cells (including image acquisition, processing, and analysis).

We selected an exogenous, non-genetically encoded dye since it facilitates rapid and inert barcoding. The most robust staining was achieved by use of fluorescent Wheat Germ Agglutinin (WGA), which selectively binds to N-acetyl-D-glucosamine and sialic acid present on a broad variety of mammalian cell types (Fig. 1B). Fluorescent WGA-conjugates are commercially available in a variety of colors with the option for customization.

We optimized the protocol for HeLa cells to obtain bright and uniform cell-surface staining, both important for image processing and barcode reading (Fig. 1B). The high intensity of the signal alongside the variety of available color-conjugates allowed us to find four colors that are easily discriminated by a common LM setup (Fig. 1B). These colors can now be mixed in various combinations: single, double, triple, or quadruple staining to obtain 15 label options (Fig. 1C).

We titrated the dye concentrations so that fluorescence intensities were comparable across all four channels, enabling reliable assessment of fluorescence profiles and accurate label assignments (Fig. 1D and Supplementary Fig. S1). Using these concentrations, we verified that the bleed-through between channels was minimal and below the levels that would be expected to hamper accurate barcode assignment (Fig. 1B, D).

Importantly, we quantified the extent of dye exchange following sample pooling and found that a maximum of 2% of cells displayed dual staining after prolonged incubation (depending on the dye combination) (Fig. 1E). Since the times measured reflect an upper limit, which can even be reduced by minimizing the delays during sample preparation, we conclude that dye exchange would not lead to gross misidentification of barcodes and therefore does not dramatically impact phenotypic profiling.

In summary, using WGA conjugated to fluorescent dyes enabled us to rapidly, efficiently, and accurately create barcodes for multiplexing of human cell samples and lay the foundation for hMultiCLEM.

### An image analysis pipeline enables barcode-decoding of human EM sample pools

Performing hMultiCLEM starts with harvesting cell populations and staining them individually using the fluorescent WGA-conjugates. Pooled samples are rapidly subjected to a well-established in-resin CLEM protocol [49] (Fig. 2A). Cryofixation by high-pressure freezing (HPF) is then utilized to avoid chemical fixation and improve preservation of cellular ultrastructure. Vitrification and arrest of cellular activity is followed by freeze-substitution and embedding in methacrylate resin that preserves dye fluorescence. Subsequently, samples are sectioned and placed on grids for transmission EM (TEM). Samples are first imaged by LM to obtain an optimal barcode reading (Fig. 2A, 2B, left) and only then subjected to contrast staining for TEM. At this step, a medium magnification EM image montage, covering the complete section area, is taken for correlation purposes only (Fig. 2A, 2B, center).

**Fig. 2:**
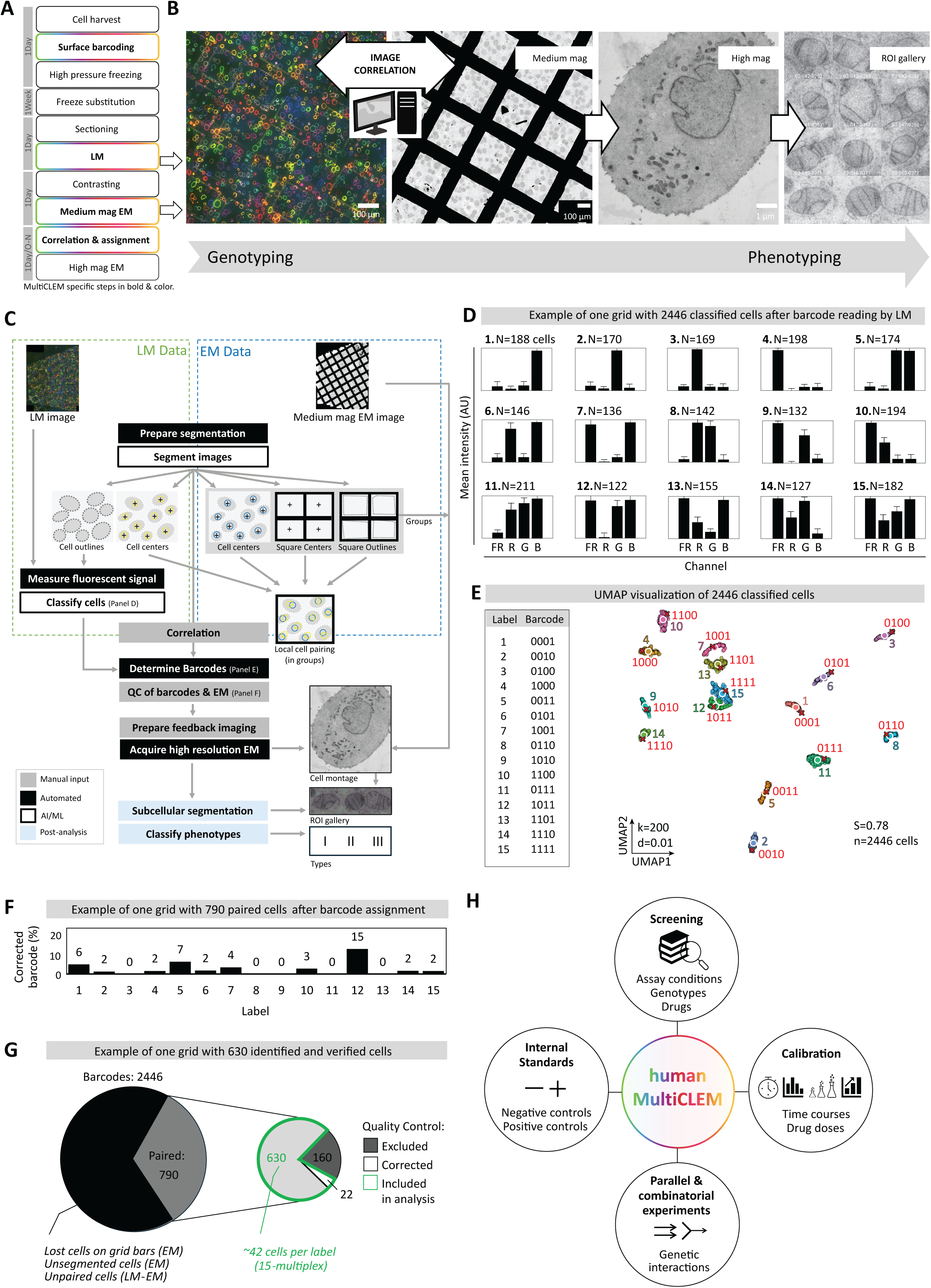
An image analysis pipeline enables barcode-decoding of human EM sample pools. HMultiCLEM includes CLEM sample preparation of multiplexed cell pellets and 3-step imaging with computational processing. **A)** Experimental workflow of the in-resin CLEM protocol designed for processing multiplexed cell pellets: Time estimates for the experimental steps are provided. **B)** Imaging workflow from genotyping to phenotyping: the workflow includes one LM and two TEM imaging steps (each automated), as exemplified. First, two corresponding images are acquired from a single EM grid for subsequent correlation (as in C). For phenotyping, high-resolution EM micrographs are acquired only at grid positions of cells successfully identified by hMClaimer. Left: Widefield LM montage, composed of four merged fluorescent channels, covering the entire sample section and showing HeLa cells labeled with 15 distinct barcodes. Center: Corresponding EM montage at medium magnification. Right: A montage of a whole HeLa cell at high magnification is shown. Galleries are generated for the regions of interest (ROI) using ImageJ. **C)** An established software workflow, hMClaimer, enables hMultiCLEM: the package includes digital image correlation and cell pairing to assign identities via barcodes, following fluorescent signal quantification and classification. Cell coordinates for feedback imaging (EM phenotyping) are determined for identified cells passing quality control (QC) (as in F). The workflow is designed to enhance accuracy and includes Artificial intelligence-driven or Machine learning-based (AI/ML) steps. **(D-G)** Computational processing is exemplified on a single grid. **D)** Barcode reading by hMClaimer: Quantification of fluorescence intensity in the EM sample is shown as mean in arbitrary units (AU). Cells are automatically assigned to one of 15 classes (labels) by K-means clustering on the normalized four-channel fluorescence intensity. N: number of cells. Dye/emission channels are denoted as B (Blue), G (Green), R (Red), or FR (Far-red). **E)** UMAP visualization of barcode reading by hMClaimer: The plot was generated with k=200 neighbors and a minimal distance (min_dist) of 0.01, showing the embedding of 2,446 cells (n, as in D) based on their normalized four-channel fluorescence intensity. Cells are colored by their class assignment (label), identified using K-means clustering on the UMAP coordinates. Large, numbered circles indicate the centroids of each cluster. Red marks show the position of the theoretical ground-truth barcodes. Proximity suggests correct cluster assignment. Cluster separation and accuracy are verified by the high average Silhouette score (S). **F)** For manual QC of barcodes prior to EM phenotyping, hMClaimer includes a cell browser with optional barcode correction or elimination of cells (due to uncertain barcodes, poor quality or localization to grid bars). Corrected barcodes (as in D) are shown as percentage per label. Note that grid-to-grid variability can occur. **G)** An example of the throughput summary: nested pie charts based on the number of cells analyzed and retained at each processing step. Discrepancies between LM and EM, for example due to limited visibility of cells on EM grid bars, are well-accepted by the high imaging speed in LM. **H)** hMultiCLEM is useful for diverse experimental set-ups, as indicated. LM: light microscopy; EM: electron microscopy; Mag: magnification; QC: Quality control.

For computational correlation of LM and EM images, essential for connecting genotype to phenotype (Fig. 2B), we established a dedicated software workflow – hMultiClaimer (hMC) (Fig. 2C). hMC provides a semi-automated platform that facilitates image correlation, assignment of cell identity, and quality control (QC). It also generates coordinates for the high-resolution TEM imaging at grid positions of only successfully assigned cells thus saving on scanning time (Fig. 2B, right). Using this platform, users can easily collect 50 cells per label for further analysis and generation of high-resolution image galleries of the region of interest (ROI) from a single EM grid (Fig. 2B, right; and for an enlarged view, see Supplementary Fig. 2). hMC can be freely accessed [50].

To verify that our WGA-based barcoding can withstand the procedure of EM sample preparation, we subjected 15 samples labeled using every possible color combination (four single, six double, four triple, and one quadruple) to hMultiCLEM. LM imaging validated that the WGA staining remains easily discernable and quantifiable (Fig. 2D). Tracking the intensity pattern that defines each barcode allows the high-confidence discrimination and assignment of all 15 color combinations that create labels for correlated cells (Fig. 2D, 2E). Still, we found that for some barcodes, the error rate can be up to 15%. To eliminate those, we introduced a visual QC interface that allows quick identification and correction of automated assignments (Fig. 2F). This step does not significantly increase the length or labor intensiveness of the protocol.

Next, we evaluated if our workflow permits collecting enough data from one multiplexed sample. The inherent limitations of EM, like large area of grid bars and inconsistent cell contrasting led to a significant number of cells that cannot be analyzed (Fig. 2G). This shortcoming is effectively compensated for by the large field of cells available for imaging on each grid (about 2000 per LM image of an entire section). After correlation, and selection of only the well-preserved cells with confident barcodes by QC, hundreds of cells (tens of cells per barcode) are still available for high resolution EM analysis (Fig. 2G).

In summary, the combination of barcoding and decoding applied in hMultiCLEM creates a platform that maximizes the information gained from each EM run. Moreover, it transforms EM into a more reliable comparative technique that minimizes inherent variation between runs by integrating internal controls with assayed samples. Together these developments allow EM imaging to be used for previously inaccessible applications: chemical and genetic screening, combinatorial experiments such as genetic interactions, time courses, dose-dependency assays, and kinetic measurements (Fig. 2H).

### hMultiCLEM enables siRNA screening

hMultiCLEM is a versatile tool that opens access to diverse experimental possibilities. We chose to demonstrate its capacity by performing genetic screening. We focused on a conserved process and cellular ultrastructure feature, the formation of cristae in mitochondria (Fig. 3A).

**Fig. 3:**
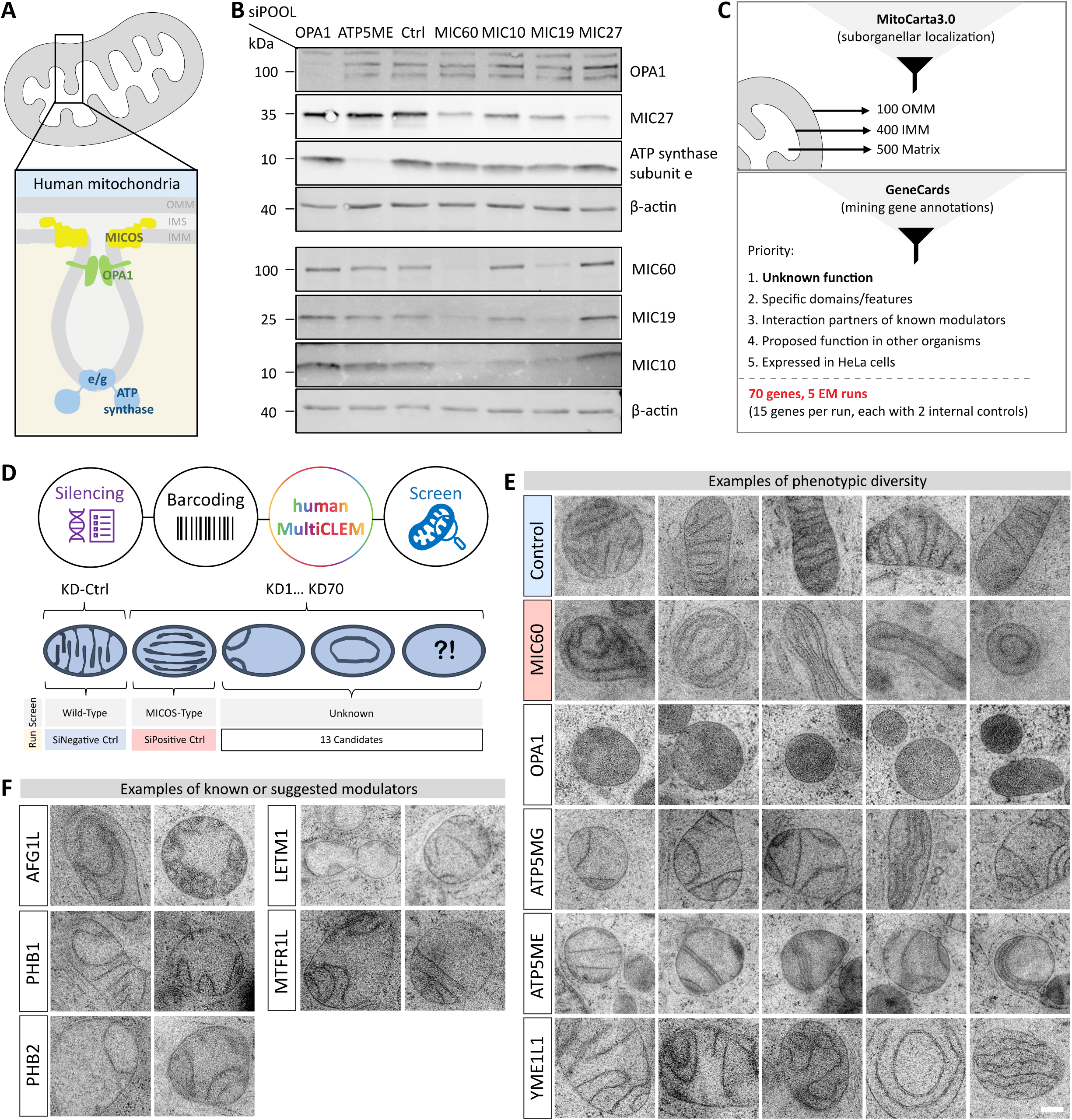
HMultiCLEM enables siRNA screening. **A)** Schematic of human mitochondrial ultrastructure: A focus on the cristae emphasizes the three core protein complexes involved in cristae organization (box). **B)** SiRNA treatment for 72 hours using high-complexity pools of 30 siRNAs (siPOOLs) is efficient for protein depletion: validated by western blotting of representative core components. Parallel depletion of non-targeted components typically results from complex instability or co-regulation. Loading control: β-actin. For a depletion kinetic see Supplementary Fig. S3A. **C)** A list of 70 candidates for screening: generated following database research and prioritization of the indicated criteria. For gene expression in HeLa see Supplementary Fig. S3B. **D)** RNAi combined with the hMultiCLEM module enables screening for unknown cristae effectors: Schemes show typical cristae phenotypes, including intact and aberrant structures in wild-type and MICOS-deficient cells, respectively. Each individual run contained a negative and positive control (Ctrl), as well as 13 candidates. KD: knock-down. **E-F)** hMultiCLEM has the capacity to capture cristae phenotypes: validated on the three core protein complexes and additional factors. Upon 72 hours siRNA treatment (as indicated), HeLa cells were subjected to hMultiCLEM. Representative micrographs show mitochondria derived from 20-50 cells. Scale bar: 100 nm. For selected high-resolution images, see Supplementary Fig. S9. **E)** hMultiCLEM captures the typical mix of phenotypes upon depletion of a single protein: exemplified for the indicated candidates. **F)** hMultiCLEM robustly detects known or suggested modulators: shown for the indicated phenotypes. OMM and IMM: Outer and inner mitochondrial membranes. IMS: Intermembrane space. ATP synthase subunits e/g and ATP5ME/ATP5MG: dimerization subunits of the ATP synthase.

Aiming at screening for additional modulators of mitochondrial ultrastructure, we employed high complexity siRNA pools (siPOOLs), each targeting a single mitochondrial protein-encoding gene. These pools are highly efficient at reducing protein levels (Fig. 3B), while keeping off-target effects very low [51]. To determine the optimal time point for the siRNA screen, we first assessed depletion kinetics by silencing transcripts of known cristae modulators for 48, 72, or 96 hours, followed by analysis of protein levels (Supplementary Fig. S3A). Though depletion efficiency for the subset of proteins tested was already very high at 48 hours, we selected 72 hours to cover the variety of proteins with differing half-lives.

Maximizing the impact of our screen, we focused on 70 samples (five EM runs with internal controls), prioritizing genes encoding mitochondrial proteins of poorly characterized or unknown function (Fig. 3C and Supplementary table S1). We enriched the selection for proteins containing domains associated with membrane binding or bending (such as BAR (Bin/Amphiphysin/Rvs) domains [52,53], coiled-coil domains [54], or amphipathic helices [55]), as well as for proteins interacting with known modulators [56], based on manual literature research and mining of public databases (NCBI Entrez Gene, UniProt). We also included proteins previously suggested to affect cristae morphology [57–62]. Importantly, we verified that our candidates are expressed in HeLa cells (Supplementary Fig. S3B).

The power of hMultiCLEM allows pre-screening for optimal conditions and controls. For example, to select the optimal media for the screen we assayed cristae morphology of untreated HeLa cells in seven different conditions (Supplementary Fig. S3C). In the absence of dramatic effects, we chose to screen in high glucose media supplemented with pyruvate and glutamine to support overall cellular metabolism and allow the survival of silenced cells with cristae alterations.

Next, we used hMultiCLEM to select a positive control with perturbed cristae, to be included in each sample. We depleted key subunits of the MICOS complex, OPA1 and the dimerization subunits (e and f) of the ATP synthase [63], which displayed their characteristic phenotypes (Fig. 3D, E). Among those, we selected the siRNA against MIC60 as the positive control due to its highly reliable and prominent phenotype.

Once controls and conditions were optimized, we performed the hMultiCLEM pipeline (Fig. 3D) with one negative control (non-targeting siRNA pool) and the positive control (siMIC60) as well as 13 candidates in each run. We analyzed 20-50 cells per label. As each cell contains multiple mitochondria (~10), we collected at least 200 mitochondria cross-section images for each sample allowing us to gain in depth insight into cristae alterations displayed (For an example gallery, see Supplementary Fig. S2).

We found variations in cristae architecture between mitochondria not only in different cells, but also within the same cell, in both control cells and upon knock-down (KD) of known modulators (Fig. 3E). While these variations could arise from biological causes (functional subpopulations, intermediate states of cristae formation, aged or dysfunctional mitochondria, etc.), or technical reasons (sectioning orientation, silencing efficiency), they are in-line with previous in-depth characterizations of MICOS which showed that depletion of its components indeed gives rise to heterogenous phenotypes [64]. Our data show that this phenotypic diversity also extends to other proteins such as the protease YME1L1 which acts on multiple substrates, including OPA1 and MIC60 [35,65] (Fig. 3E). Importantly, capturing such diversity and its meaningful biological variation underscores the value of a large sample size, as provided by hMultiCLEM.

hMultiCLEM allowed us to rapidly ascertain cristae phenotypes upon KD of known or suggested modulators (Fig. 3F). For example: the multifunctional regulators Prohibitin 1 and 2 [66,67], as well as the AAA-ATPase AFG1L/LACE1, which interacts with YME1L1 and is implicated in protein homeostasis and OXPHOS regulation [62]. We also identified phenotypes for two newly suggested cristae modulators published during the course of our screen: Letm1, initially described as an IMM ion transporter and recently shown to sculpt membranes in liposome assays [43]; and MTFR1L, a regulator of mitochondrial dynamics [68].

More generally, the ability of hMultiCLEM to rapidly reproduce known phenotypes proves the capacity of our screening approach to compare multiple distinct suggested modulators in a single coherent study.

### Systematic analysis of top candidates explores diverse functional mechanisms

Our ability to assign phenotypes to known or newly suggested cristae modulators encouraged us to select candidates exhibiting striking phenotypes that had not previously been implicated in cristae architecture regulation (at the time of the study) for further analysis. We selected eleven top candidates (hits) that either localized to the OMM, IMS, or were associated with the IMM based on MitoCarta3.0 [69] (Table 1). Among the hits of unknown function, a prevalent phenotype was a reduction in cristae number (Fig. 4A), suggesting a fundamental role in cristae formation. Interestingly, many of the hits chosen for follow-up seemed connected to lipid homeostasis [61,70–73]. Because of shared phenotypic similarities (for example waved cristae stacks, Fig. 4A), we also included SLC25A46, implicated in Leigh Syndrome, which is known to affect cristae architecture potentially through regulating lipid transfer [61,74].

**Fig. 4:**
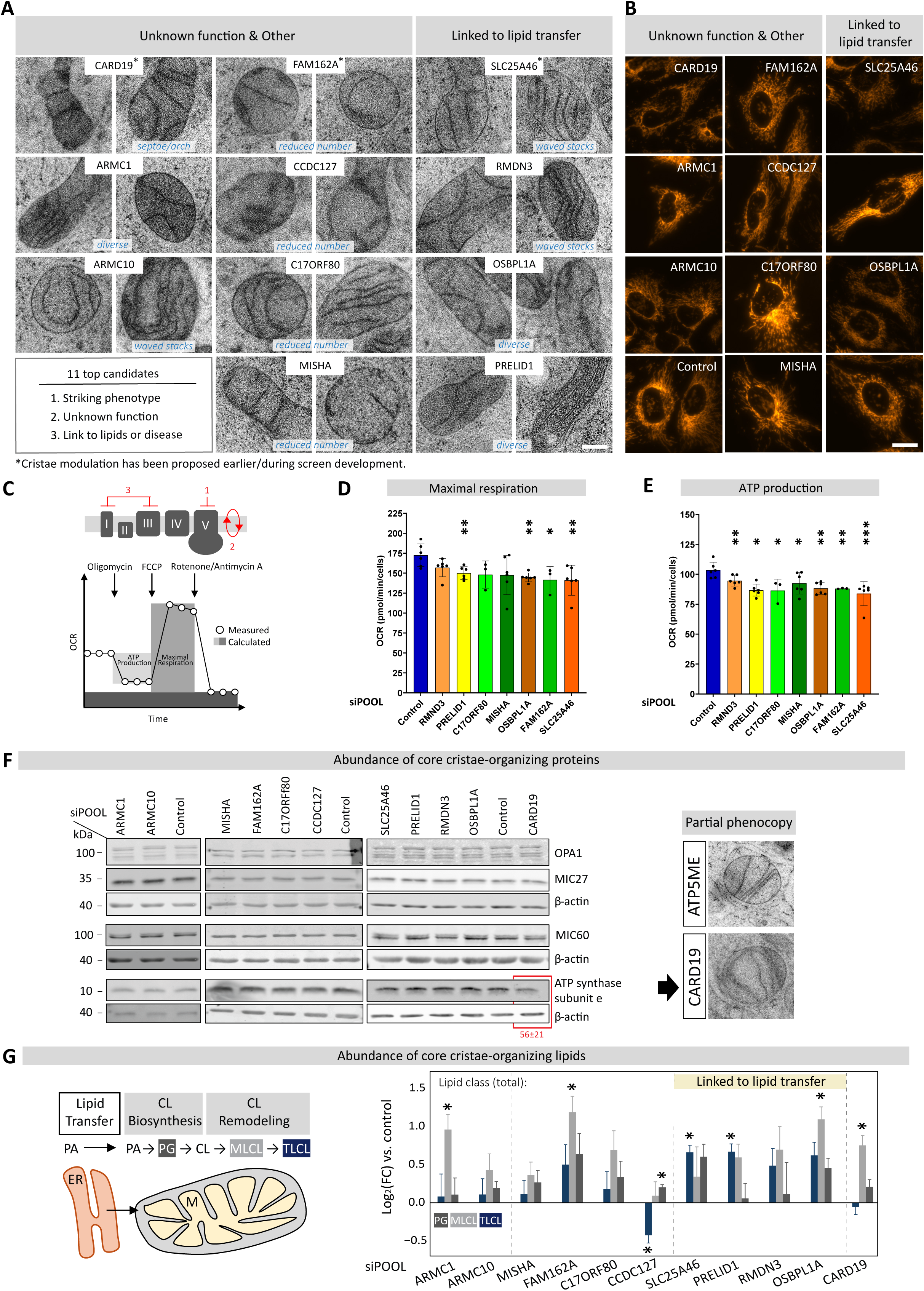
Systematic analysis of top candidates explores diverse functional mechanisms. The top candidates of the hMultiCLEM screen were selected based on their ultrastructural phenotype according to the depicted criteria: Upon 72 hours siRNA treatment (as indicated), HeLa cells were subjected to diverse follow-up analyses. **A)** Cristae morphology, as revealed by hMultiCLEM, is shown for each of the 11 top candidates by two representative micrographs of mitochondria. For an example gallery and repeats see Supplementary Figs. S2 and S4, respectively, and for selected high-resolution images, see Supplementary Fig. S9. Scale bar: 100 nm. **B)** Mitochondrial network architecture was assessed by live cell staining with MitoTracker Orange followed by automatic imaging. Representative images of cells are shown. Scale bar: 20 μm. **C-E)** Respiratory capacity was assessed by Seahorse measurements of oxygen consumption rates (OCR) on whole cells. **C)** Scheme. **D)** Maximal respiration rate and **E)** ATP production are shown for candidate proteins demonstrating significant effects. Data represent mean ± SD and were analyzed using an unpaired two-tailed Student’s t-test (relative to control, n=3-6). *p < 0.05; **p < 0.01; ***p < 0.001. For OCR profiles, see Supplementary Fig. S5. **F)** Levels of cristae-organizing proteins assessed by immunoblotting: antibodies against the three core players: MICOS (represented by MIC60 and MIC27, as central and lipid-associated components, respectively), Opa1 and ATP synthase subunit e were used. Loading control: β-actin. Representative blots are shown. For quantifications of 3 to 6 repeats, see Supplementary Fig. S6A. **G)** Levels of Cardiolipin and pathway intermediates from lipidomic analysis on whole cells: Data were normalized to the total protein content. Analysis of total lipid classes for mature cardiolipin (CL), monolysocardiolipin (MLCL) or phosphatidylglycerol (PG) are shown (scheme). Bar graphs represent the log₂ fold change (Log₂FC, calculated from means of 4 repeats) of indicated mitochondrial lipid classes relative to control cells. Data were analyzed using an unpaired two-tailed Welch’s t-test. *p < 0.05. KD: Knock-down. Ctrl: Control. For analysis of individual lipid species see Supplementary Fig. S6B.

**Table 1:** Top hits of hMultiCLEM screen on mitochondrial ultrastructure. Cristae phenotypes described previously/during the course of the screen are highlighted alongside the year of publication (bold/italics), whereas links to disease and organismal level are highlighted in bold.

| Gene<br>(Uniprot ID) | Localization/<br>Association | Literature | Our study | Conclusion/<br>Hypothesized mechanism |
| --- | --- | --- | --- | --- |
| **CARD19**<br>(Q96LW7) | OMM/<br>MICOS [44] | ***Moderate impact on cristae in macrophages and fibroblasts (2022)*** [44]<br>Redox sensor [107]<br>**Immune response [106, 144]** | Robust and strong impact on cristae<br>Respiration defect<br>Lowered ATP synthase e levels | OMM-IMM crosstalk<br>Effector of ATP synthase |
| **ARMC1**<br>(Q9NVT9) | OMM/<br>MICOS [76] | Mitochondrial distribution and dynamics [76, 130]<br>No cristae defect upon knockout [76] | Mild cristae defect<br>MLCL accumulation | Cristae alterations may represent adaptive changes aimed at restoring mitochondrial function ('bagpipe hypothesis' [133]) |
| **ARMC10**<br>(Q8N2F6) | OMM | Mitochondrial distribution and dynamics [130, 131]<br>Energy-linked stress [132] | Cristae defects in a subset of mitochondria | Possible context-dependent function |
| **SLC25A46**<br>(Q9H4I9) | OMM/<br>OPA1 [61]<br>EMC [134]<br>MIC19 [134] | ***Leigh syndrome with cristae defect (2016)*** [61]<br>Mitochondrial dynamics [74]<br>Regulation of lipid influx [61]<br>**Neurodegeneration [143]** | Cristae defects<br>Respiration defect<br>Mitochondrial fragmentation | Signature of lipid-linked cristae effector: wavy stacks, septae, arches |
| **RMDN3/PTPIP51**<br>(Q9Y2H9) | Mitochondria-ER contact | Transfer of PA, MLCL and lipid radicals [72, 135]<br>**Neurodegeneration [136]** | Discrete cristae alterations<br>Respiration defect<br>No change in levels or assembly of core proteins | Cristae defect potentially linked to lipid transfer, stress or disrupted contacts |
| **PRELID1**<br>(Q9BT81) | IMS/<br>OPA1 [70] | PA transfer OMM to IMM [70, 71, 75, 147]<br>Mitochondrial dynamics [75, 41, 147]<br>Membrane binding/mild bending [137] | Diverse cristae alterations<br>Respiration defect<br>Mitochondrial fragmentation | May stabilize membrane curvature beyond lipid transfer |
| **OSBPL1A/**<br>**ORP1L**<br>(Q9BXB5) | Cytosol;<br>Mitochondria-ER-lysosome<br>three-way<br>contact [138] | Cholesterol transfer,<br>lipid sensor [142] | Cristae defect in subset of<br>mitochondria<br>Respiratory defect<br>Lipid imbalance | Likely broad impact on mitochondrial<br>function, possibly via disruption of<br>contacts |
| **CCDC127**<br>(Q8N6U8) | OMM;<br>Mitochondria-ER contact [92] | Contact regulation,<br>lipid droplet homeostasis,<br>mitochondrial dynamics [139]<br>**Adipocyte differentiation [139]** | Reduced number of cristae<br>Respiration defect<br>Reduced CL levels | Cristae defect potentially linked to<br>lipid transfer and/or disrupted<br>contacts<br>CL deficiency and high expression in<br>tissues suggests link to disease |
| **FAM162A**<br>(Q9NVV4) | Cristae [140] | **Cristae defect in COS7 cells (2024)**<br>**[45, preprint]**<br>Mitochondrial dynamics [45]<br>Promotes longevity and apoptosis<br>under hypoxia [45]<br>Reduced OPA1 levels upon KD [45]<br>**Overexpressed in cancers [45]** | Reduced number of cristae<br>Fragmented, rounded mitochondria<br>Respiration defect<br>No change in OPA1 levels or assembly | Broad tissue expression suggests<br>fundamental role mitochondrial<br>function |
| **C17ORF80**<br>(Q96HY7) | Matrix:IMM/<br>mtDNA<br>[93, 141] | Potential mtDNA maintenance [141]<br>Unclear functional impact on mtDNA-binding [93]<br>Potentially stress-responsive [93] | Reduced number of cristae<br>Mitochondrial network alterations<br>Respiration defect<br>No impact on ATP synthase e levels or<br>assembly<br>Low expression across tissues | Potential tissue- and condition-<br>specific function<br>Potential functional replacement or<br>sequestering of ATP synthase subunit<br>f by a predicted homology domain<br>[93] |
| **FAM136A/**<br>**MISHA**<br>(Q96C01) | IMS/<br>MIC19 [119] | Mitochondrial stress [115]<br>Proteostasis (specific proteins and<br>tissues) [96, 116]<br>**Ménière's disease [100]** | Reduced number of cristae<br>Respiration defect<br>Unique (co-)expression pattern with<br>disease relevance<br>Oligomerization | Proposal of an unrecognized<br>mechanism for initiating cristae<br>formation through membrane<br>protrusions, stabilized by ring-like<br>structures with potential disease<br>relevance |

OMM/IMM: Outer/Inner mitochondrial membrane; IMS: Intermembrane space; CL: Cardiolipin; MLCCL: Mono-lyso cardiolipin; PA: Phosphatidic acid; ER: Endoplasmic Reticulum; EMC: Endoplasmic reticulum Membrane protein Complex, acting in protein insertion and lipid regulation, [145, 146];

We first re-imaged the hits by hMultiCLEM and verified that their phenotypes are reproducible in a small-scale secondary screen (Supplementary Fig. S4). Next, we assessed the effect of protein depletion on gross mitochondrial morphology using LM imaging of MitoTracker-stained cells. We found that depletion of the lipid modulators SLC25A46 and PRELID1 resulted in fragmented networks, as described previously [74,75]. Mitochondrial fragmentation was also observed upon KD of uncharacterized FAM162A. A very pronounced phenotype, with mitochondria appearing as scattered patches, was displayed by cells silenced for C17ORF80 (Fig. 4B).

To determine whether the altered cristae displayed by our hits have functional relevance for respiration, we measured the oxygen consumption rates (OCR) of whole cells (Fig. 4C and Supplementary Fig. S5). Focusing on calculations of maximal respiration (Fig. 4D) and ATP production (Fig. 4E), we found a reproducible reduction in all hits, except ARMC1 and ARMC10 (Supplementary Fig. S5). Notably, the impact we observed on cristae was comparatively minor in ARMC1, and was not detected in a previous knockout study [76]. Regardless, this finding demonstrates that hMultiCLEM effectively captures subtle and complex phenotypes.

A change in cristae morphology could arise for a variety of reasons – indirect effects by regulating known modulators and lipid composition or by novel, unexpected pathways. To explore the underlying causes and organize the new effectors into functional categories, we first assayed their effect on the abundance of the main cristae modulators – the MICOS complex (represented by MIC60 and MIC27, as central and lipid-associated components [30,77,78], respectively), OPA1, and ATP synthase subunit e. Surprisingly, we found that most of our new effectors do not work by regulating the levels of these proteins. However, there was one interesting exception: KD of Card19 caused a ~44% reduction in ATP synthase subunit e protein levels (Fig. 4F and Supplementary Fig. S6A). Of note is that the septa- and arch-like cristae traversing and dividing the matrix observed in Card19 KD cells (Fig. 4F, right, and Fig. 3E) phenomimick ATP synthase dimerization defects [20,28,79]. The finding that loss of Card19 partially phenocopies loss of ATP synthase subunit e suggests that it affects cristae by regulating, directly or indirectly, the ATP synthase assembly.

Since most of the hits did not seem to affect the core proteins forming cristae, we hypothesized that some may impact cristae through regulation of membrane composition [19,31,78,80]. To this end, we measured levels of mature cardiolipin (tetralinoleoyl-cardiolipin, TLCL, or from here on CL), the intermediate mono-lyso-cardiolipin (MLCL) and their immediate biosynthetic precursor phosphatidylglycerol (PG) (Fig. 4G, left) using whole-cell lipidomics [81]. We found that KD of four proteins lowered individual CL species (CCDC127, C17ORF80, CARD19 and ARMC10, Supplementary Fig. S6B), and KD of CCDC127 substantially reduced the total amount of mature CL by ~30% (Fig. 4G). Reduction of CL, associated with numerous pathologies [31,82], is of clinical importance and suggests one mechanism by which CCDC127 impacts mitochondrial ultrastructure. Surprisingly, silencing of the lipid transport-associated proteins (highlighted in yellow, Fig. 4G), and FAM162A, elevated the levels of CL, MLCL, and PG to various extents, which may underlie or arise from their individual cristae phenotypes [83,84], or result from compensatory effects (Fig. 4G). Similarly, depletion of CARD19, ARMC1, and C17ORF80 elevated MLCL levels, a typical consequence of CL remodeling defects or degradation. CL degradation also occurs in the context of ATP synthase dimerization defects [83,84]), and serves as a marker of mitochondrial stress, which may contribute to their ultrastructural phenotypes.

In summary, our secondary assays support the validity of our initial hit assignment and, moreover, raise hypotheses regarding the mechanisms of action of several candidates. However, the majority have an effect that is independent of the known proteins and lipids, suggesting exciting new avenues for studying cristae formation and regulation.

### Expression profiles suggest functional networks and their tissue-specific relevance

An interesting aspect of our secondary screens is that most hits do not seem to impact the known pathways for cristae formation. To gain insight into their phenotype, we used an unbiased approach by assessing the protein abundance profiles across human tissues and organs measured by mass spectrometry [85] to identify co-expression patterns. First, we examined the abundance of MICOS components across various tissues. Notably, while MICOS components are ubiquitously expressed, we could observe divergent expression patterns for the two functional subcomplexes of MIC60 and MIC10 (Supplementary Fig. S7A). Since the two subcomplexes are thought to contribute differently to cristae organization [64,86], the divergence of their expression patterns suggests that a certain fraction of cristae morphology at the tissue level may arise from different subcomplex abundances.

Following the hypothesis that proteins highly expressed in a given tissue help shape its characteristic mitochondrial architecture (Fig. 5A), we focused in on the expression levels of our hits. We observed greater variation among the hits compared to MICOS components (Fig. 5B). Except for FAM162A, which is highly expressed across most tissues, the remaining candidates exhibited distinct tissue-specific expression patterns. For example, brain tissues showed higher abundances of ARMC10 and RMDN3 or OSBPL1A and FAM136A/MISHA, respectively. In contrast, the liver exhibited high levels of RMDN3, whereas the heart was enriched with MISHA. When we explored correlations between protein abundance and ultrastructure, we noted that liver cells (that exhibit higher RMDN3 levels) tend to have sparse crista. Indeed, RMDN3-KD resulted in more organized cristae (Fig. 4A). Conversely, cerebral cortex and heart cells (that display high MISHA levels) tend to have dense cristae [87] correlating well with the phenotype of MISHA depletion showing a marked reduction in cristae (Fig. 4A). Whether these proteins indeed contribute to cristae morphology in the respective tissues requires additional exploration and represents an important direction for future work

**Fig. 5:**
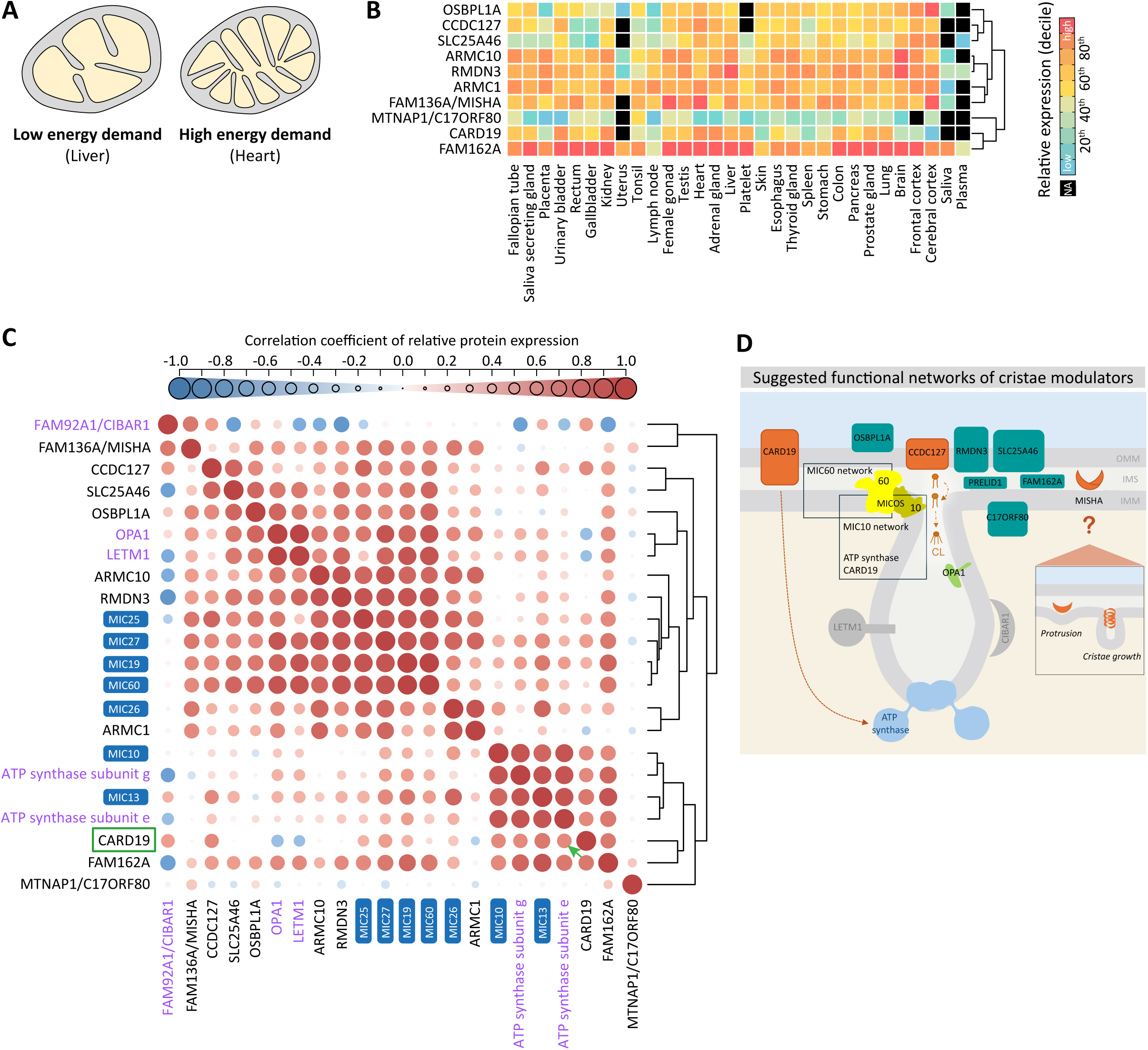
Expression profiles suggest functional networks and their tissue specific relevance. **A)** Illustration of mitochondria types with ultrastructural differences related to different cellular energy demands. **B)** Protein levels across human tissues for the top screening hits. Abundances were extracted from the integrated human tissue datasets in PaxDB 6 and converted to per-tissue percentiles following normalization. Tissues were hierarchically clustered based on Pearson correlation of their quantified proteomes. PRELID1 is not shown due to the limited available data. For core proteins, see Supplementary Fig. S7A. NA: data not available in datasets. **C)** Co-expression and correlation analysis: performed on a curated subset of proteins, including the top screening hits, the three core protein complexes (MICOS subunits, OPA1, ATP synthase subunits e/g), and the newly suggested membrane-shaping proteins LETM1 and CIBAR1. A Pearson correlation matrix based on tissue expression percentiles (as in A and Supplementary Fig. S7B) was used. Large blue circles indicate antagonistic expression, and large red circles show strong co-expression. Hierarchical clustering was used to identify groups pointing to potential functional networks, for example CARD19 and ATP synthase subunit e (green, see also Fig. 4F). Purple: Membrane-shaping proteins (defined by membrane-bending activity on model membranes). **D)** Schematic of the top hits identified in the screen (turquoise and orange) integrated into the current framework of cristae organizing proteins (gray: additional membrane shaping proteins): Functional networks are proposed as indicated (boxes). In three cases (orange), a potential mechanism for impacting cristae is suggested in this work. The insert depicts the direct membrane-shaping activity hypothesized for MISHA.

We then explored correlations between the protein abundances of known cristae modulators alongside our hits, to identify potential functional cooperations based on co-expression patterns. Pearson correlation analysis revealed distinct groups with similar abundance profiles across different tissues (Fig. 5C). The largest cluster contains the MIC60 subcomplex which is distinct from the second major cluster comprising MIC10 and MIC13 of the MIC10 subcomplex, supporting the notion of unique roles of these two subcomplexes [64,86]. The independence of MIC27 and MIC26 from the MIC10 cluster was unexpected and merits further investigation. Proteins with membrane-shaping activity demonstrated on model membranes [18,26–28,43,88–90] are found both within MIC10 and MIC60 clusters, and outside (highlighted in purple, Fig. 5C). Interestingly, CARD19 (green rectangle, Fig. 5C) is part of the MIC10 cluster alongside the ATP synthase e/g subunits, supporting our observation of CARD19 impacting ATP synthase e protein levels (Immunoblot, Fig. 4F), as well as the proposed crosstalk between the MICOS and ATP synthase machineries [86,91]. Moreover, proteins with demonstrated physical interactions with MIC60 or its subcomplex members, such as CCDC127 [92] (preprint) and SLC25A46 [61], indeed reside in the same cluster. Closer inspection reveals sub-groups within the MIC60 cluster, for example: LETM1 (recently shown to shape membranes and affect OPA1 oligomerization, [43]) indeed clusters with OPA1 (Fig. 5C, Supplementary Fig. S7B).

C17ORF80 does not cluster with any of the other proteins and, indeed, it has the lowest abundance across tissues, suggesting that it may be a tissue-specific or condition-specific modulator. Interestingly, the structural homology between C17ORF80 and ATP synthase subunit f suggests that C17ORF80 might replace or sequester subunit f in unique settings [93].

A second hit that clusters independently from the other proteins is CIBAR1/FAM92A1, recently shown to shape membranes [88]. Because CIBAR1 showed limited detection among the available tissue-based datasets, reliable analysis was challenging (Supplementary Fig. S7B). Nevertheless, the most notable subcluster comprises CIBAR1 and MISHA, the only protein co-segregating with CIBAR1. This separation from clusters containing the MICOS subcomplexes supports the idea that CIBAR1 and MISHA may act via an independent mechanism, making them interesting targets for further study. Simultaneously, in addition to clustering with CIBAR1, MISHA shows also similarity to the MIC60 cluster, without falling into a specific sub-group, unlike most other proteins.

In summary, the analysis of tissue distributions provides a foundation for a holistic view of the network of proteins modulating cristae (Fig. 5D, Table 1).

### MISHA is a Ménière’s disease-linked IMS protein impacting cristae formation with the ability to oligomerize

The secondary screens highlighted one hit, MISHA, paving the way for further mechanistic investigation. Among the most fundamental cristae phenotypes, MISHA shows broad expression and is enriched in clinically relevant tissues with a unique co-expression signature, yet does not alter the levels of MICOS, OPA1, ATP synthase and CL, suggesting an independent mode of action. Moreover, its IMS localization positions it as a potential direct membrane-effector.

To ascertain whether MISHA silencing is indeed causal for the cristae phenotypes, we first validated that the employed siRNA Pool effectively depletes MISHA (Supplementary Fig. S8A). Next, although MISHA did not affect the individual protein levels of known direct effectors, we verified that MISHA does not affect their complex assembly and oligomerization – a common theme enabling membrane sculpting. Blue native PAGE analysis clearly showed that MISHA silencing (as well as silencing of RMDN3, FAM162A, or C17ORF80) does not impact the assembly of major membrane-shaping machineries, including MICOS (represented by its core components MIC60 and MIC10), OPA1 and ATP synthase (assayed via its catalytic core subunit beta to assess complex integrity independently of monomeric, dimeric or higher assembly states) (Fig. 6A).

**Fig. 6:**
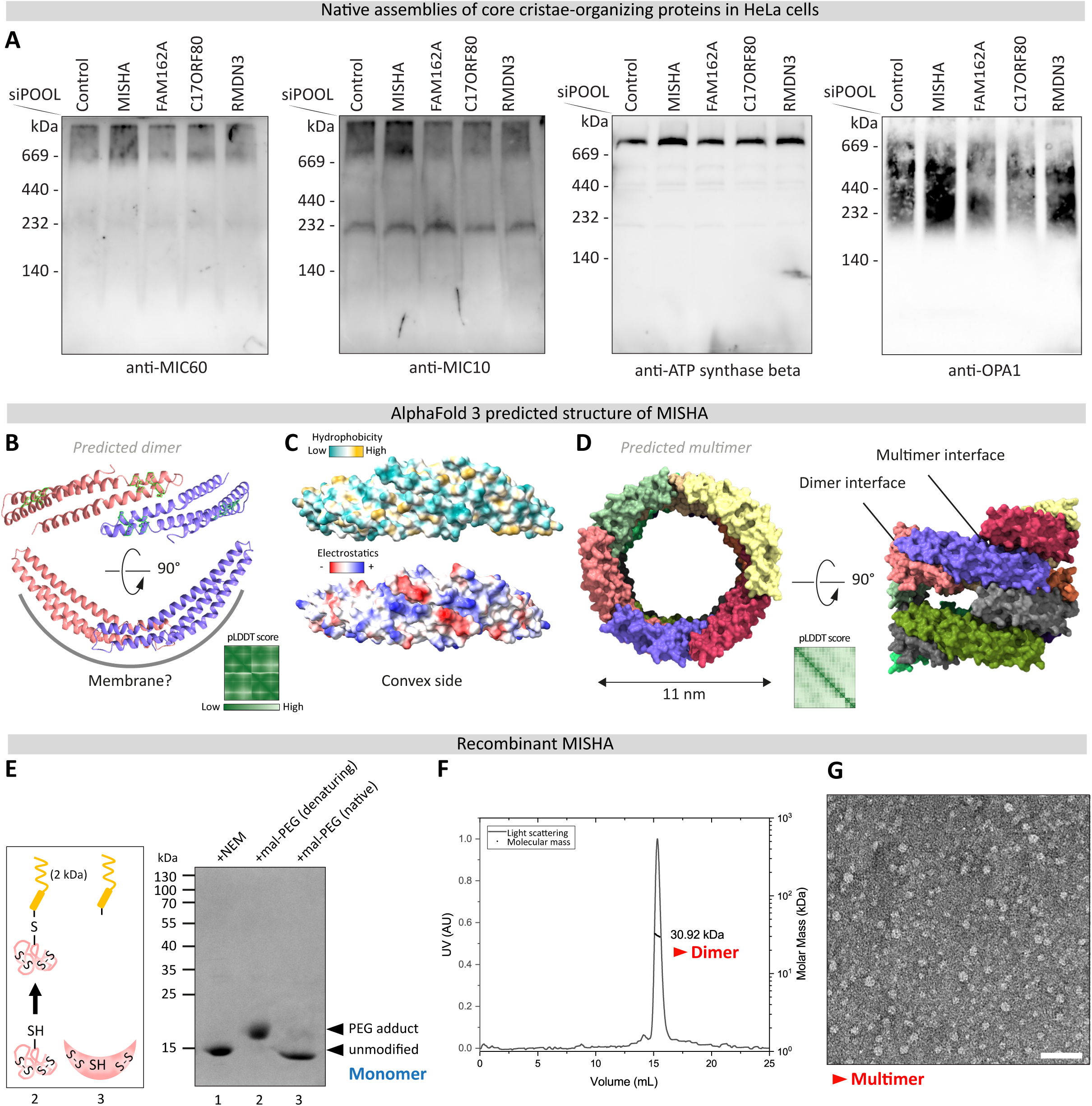
MISHA, a Ménière’s disease-linked IMS protein, forms dimers and has the capacity for self-assembly. **A)** Analysis of the native complex assembly of core proteins using Blue native PAGE: following 72 hours depletion of the indicated candidates, HeLa cells were harvested, and mitochondria-enriched fractions were immunoblotted and probed with antibodies against MIC60, MIC10, OPA1 and ATP synthase subunit β. **B-D)** AlphaFold 3 predicted structures of MISHA. **B)** Dimer, with cysteine side chains highlighted. **C)** Dimer, with surface representations. Top, hydrophobic (yellow) and charged surface (cyan). Bottom, electrostatic potential surface (negative in red; positive in blue). **D)** Prediction of MISHA higher-order assembly state. The heatmaps display pLDDT scores (predicted Local Distance Difference Test), indicating the model’s per-residue confidence (green = low, white = high). For enlarged views of scores see Supplementary Fig. S8B-C. **E)** SDS-PAGE gel of purified recombinant MISHA: showing the expected size for a monomer (for mass spectrometry, see Supplementary Fig. S8D). The thiol status by PEG-modified maleimide (mal-PEG, ~2 kDa, yellow) migration assay is consistent with the AlphaFold 3 model. Adding PEG-mal to the denaturing SDS gel-loading buffer (lane 2, and ‘2’, cartoon) caused a mobility shift (arrow, cartoon) relative to N-ethylmaleimide (lane 1, NEM, 125 Da), demonstrating one unpaired thiol (SH) that did not react in its native, folded state (lane 3, and ‘3’, cartoon), consistent with the lack of surface exposure. Accordingly, paired thiols (S-S, two disulfide bonds) exist in the MISHA monomer (light pink, cartoon). **F)** Analysis of the solution molecular mass of recombinant MISHA: determined by SEC-MALS analysis corresponds to a dimeric state. **G)** Negative-stain TEM image of recombinant MISHA demonstrating assembled structures. Scale bar: 50 nm.

How then does MISHA impact mitochondrial ultrastructure? To gain insights into a potential mechanism we modeled MISHA structure using AlphaFold 3 [94]. The predicted monomer consists of a bow-shaped three-helical bundle held together by two twin disulfide bridges leaving one free cysteine. This aligns with MISHA’s localization to the IMS, where twin-cysteine motifs commonly enable Mia40-dependent import [95], as recently confirmed for MISHA [96]. Interestingly, the predicted structure for a MISHA homo-dimer showed a banana-shaped assembly with an extensive dimer interface (Fig. 6B and Supplementary Fig. S8B; and seen also by [96]), an architecture typical of the BAR family of membrane-bending proteins [97,98]. We found hydrophobic and positively charged amino acids on the convex side, suggesting interaction with the negatively charged head-groups of the IMM (Fig. 6C). Models of a higher-order multimer revealed putative ring-shaped structures, potentially assembling into stacked or spiral architectures, with dimensions matching the internal diameter of cristae in respiration-active mitochondria (11 nm) [99] (Fig. 6D and Supplementary Fig. S8C). Together, these structural features raise the hypothesis that MISHA directly binds membranes, induces negative curvature, and promotes membrane protrusions as a potential new mechanism for generating cristae through inverted scaffolding. Further self-assembly into higher-order structures may drive growth and dictate cristae width.

To test the validity of the predicted structures, we produced recombinant MISHA (Uniprot Q96C01, human FAM136A) for biochemical analysis. We found that purified MISHA migrated as a monomer by SDS-PAGE, which is consistent with the molecular weight verified by mass-spectrometry (~15.6 kDa) (Supplementary Fig. S8D), while allowing for a non-covalent dimer. To test the disulfide bonding status, we used PEG-modified maleimide (PEG-mal), which reacts with unpaired cysteines. In this assay, migration of MISHA was retarded to an extent consistent with modification of one cysteine and eight cysteines protected in disulfide bonds as predicted by AlphaFold 3 (Fig. 6E). In support of the capacity of this purified monomer to dimerize, we found by size exclusion chromatography-multi-angle light scattering (SEC-MALS) a solution molecular mass for MISHA consistent with a dimer (~30.9 kDa measured versus 31.3 kDa calculated molecular mass) (Fig. 6F). Notably, the predicted MISHA dimer structure buries the unpaired cysteine in the dimer interface. Indeed, PEG-mal treatment prior to denaturation did not modify MISHA (Fig. 6E). Moreover, capacity for higher-order assemblies was observed by negative-stain EM, showing ring-like structures roughly matching the expected size of 11 nm (Fig. 6G), thus providing experimental evidence supporting the structure prediction.

In summary (Fig. 5D, Table 1), our data places MISHA as an IMS protein having the potential to impact cristae by a novel mechanism. This mode of action may underlie its role in the etiology of Menier’s disease [100].

## Discussion

In this study we established hMultiCLEM, a scalable and multiplexed EM platform that enables genetic screening in human cells through fluorescent barcoding. Using this approach, we performed the first systematic ultrastructure survey of mitochondrial proteins and identified several previously unrecognized cristae modulators. With the goal of integrating these proteins into existing models of cristae organization, we used biochemical and computational analyses to distinguish between direct and indirect effectors. For three proteins, our findings enabled us to suggest potential mechanisms for their impact on cristae architecture, notably through regulation of the ATP synthase (CARD19), CL levels (CCDC127), and an independent mechanism (MISHA/FAM136A) (Fig. 5D). These findings broaden the inventory of cristae modulators towards a comprehensive view and highlight the utility of hMultiCLEM for discovering ultrastructure modulators at scale.

hMultiCLEM represents a leap in the capability of EM to be efficiently utilized for screening as well as other increased throughput requirements. We foresee that hMultiCLEM will be further optimized for even higher throughput by more complex barcoding (potentially using methods other than fluorophores). hMultiCLEM gives rise to vast amounts of data supporting future efforts to automate EM image analysis, similarly to what is now being demonstrated for volume and non-correlative EM [101–103]. Developing non-fluorescent barcodes could avoid lower-contrasted correlative EM, enhance ultrastructure visibility, and thereby reduce challenges in automated analysis. Future development of powerful automated (Artificial Intelligence (AI) based) computational analysis pipelines should also enable in-depth characterization of phenotypes including clear classification into the various structural groups (Fig. 3D). Moreover, AI based analysis software should enable quantification of intra- and inter-cellular subpopulations, allowing examination of their contribution to structural heterogeneity and revealing their biological significance.

Additionally, the capacity for scanning thousands of mitochondrial EM images from various genetic backgrounds enabled the identification of rare and lesser-studied phenotypes. For example, we could detect mitochondria whose cristae are branched, looped or networked (Supplementary Fig. S9). These phenotypes resemble, and potentially extend beyond, previous characterizations linked to imbalance of MICOS and ATP synthase [22] or OPA1 isoforms [104], or to IMM QC [105]. Although these morphologies were rare, their unique occurrence in depleted cells suggests they may represent meaningful biological states arising from genetic perturbation. More images from these depletion backgrounds, across diverse conditions and cell types, are needed to further explore these structures and their causes.

While many of the proteins in our screen were already suggested to impact cristae architecture in previous reports, the fact that each lab works on a different cell type in varying conditions makes it difficult to ascertain the generality of findings. The power of hMultiCLEM is that it enables easy comparison between different genetic backgrounds in the exact same cells, conditions, and even samples. Thus, the presented hMultiCLEM dataset positions us to interpret our findings within an integrative context (as summarized in Table 1).

For example, during the development of our screen, the OMM protein CARD19 was reported to moderately affect cristae morphology in macrophages and fibroblasts, potentially through interactions with MICOS in the context of immune signaling [44,106]. Based on its crystal structure, a role as a redox sensor was proposed [107]. In our HeLa cell system, cristae abnormalities were robust and partially mimicked ATP synthase dimerization defects, which was confirmed by reduced ATP synthase e levels. Further supported by shared similarities in the expression pattern, CARD19 emerges as an effector of ATP synthase assembly or stability in multiple cell types, beyond the context of immune response.

Interestingly, several hits from our screen were linked to lipid transport from the Endoplasmic Reticulum (ER) to mitochondria, including the transfer of phosphatidic acid (PA), essential for CL biosynthesis and cristae growth [108]. Surprisingly, none led to a depletion of CL, instead, we observed differential accumulation of CL species and its biosynthetic precursor. While often reflecting defects in CL biosynthesis or remodeling, disruptions may also point to compensatory reorganization and redundancy in lipid transfer pathways between the ER and mitochondria, as we have previously shown in yeast [109,110]. Under this lipid imbalance, cristae became disorganized, highlighting both shared and distinct features of proposed lipid transfer proteins. Notably, MLCL accumulation in the IMM hampers protein interactions, due to its lower binding affinity compared to CL [33], and destabilizes complex assemblies that maintain cristae morphology. Conversely, cristae defects may exacerbate this instability, promote degradation of released CL, and further drive MLCL accumulation [84], creating a self-reinforcing cycle. Elevated CL levels may then reflect saturation or a compensatory response, in which it is either free or functionally engaged in cristae enrichment. These findings underscore the complex interplay between lipid homeostasis and cristae organization, warranting further investigation.

Of note, since many of these proteins locate to the OMM and also act as organelle contact site tethers, the fragmentation of mitochondria that we observed may indirectly result from disrupted contacts to the ER causing cellular stress and disrupted lipid transfer, as well as fission/fusion defects [111–113].

However, the most compelling finding was the poorly characterized soluble IMS protein MISHA/FAM136A, as a potential direct effector of mitochondrial ultrastructure and respiratory function. MISHA is evolutionary conserved across metazoans, and influences locomotion and behavior in *C. elegans* [114]. Several splice isoforms of transcripts are predicted, with one variant that is of primary focus, encoding a protein recently confirmed to locate to mitochondria [96,115,116]. A MISHA mutation has been identified in patients with familial Ménière’s disease [100], leading to protein truncation and failed import into mitochondria [116]. MISHA knockout (KO) mice exhibit hearing loss [117]. Examining available proteomics data, we found MISHA expression to be widespread, with particularly high levels in tissues with high energy demand, such as heart and brain. Just this year, the first cellular findings revealed multiple mitochondrial dysfunctions, i.e. elevated Reactive Oxygen Species (ROS) production, reduced membrane potential, and decreased ATP production upon MISHA-KD, consistent with our seahorse data [115]. In addition, MISHA was identified in a screen for genes essential for respiration and was suggested to play a role of a chaperone in supporting IMS protein homeostasis, with its loss triggering the mitochondrial integrated stress response (mtISR) [96,116]. The extensive study encompassed diverse KO cells, derived from different mouse organs and Ménière’s patients demonstrating that the effects on protein levels are tissue specific. In our HeLa cell system, we did not observe a notable effect on protein abundance and complex assembly for the MICOS subcomplexes, OPA1, or ATP synthase. We further did not observe an alteration in OPA1 processing that would indicate an activation of the mtISR under the assessed conditions. It may be that the protein instability described in the KO represents a tissue-specific activity or a later consequence, or compensatory effect, of the MISHA-induced ultrastructure defect. Ménière’s disease is emerging as a potential mitochondrial disorder with MISHA implicated in the underlying disease mechanism, making it an interesting therapeutic target. Regardless, its pronounced expression in heart tissues, with a cristae width matching the predicted MISHA ring diameter, highlights additional research directions with strong relevance to mitochondrial pathology.

What is the mechanism of action for MISHA? Our work did not find indications for effects of MISHA KD on the three key players in cristae formation, nor on levels of CL. Therefore, we used AlphaFold 3 structure predictions to generate a hypothesis for MISHA function. An intriguing dimeric state of MISHA, which we confirmed by SEC-MALS using recombinant protein, strikingly also exists in cells, as has been shown recently [96]. In addition, we found evidence for previously unrecognized higher-order assemblies of MISHA, including the prediction of stacked or spiralling ring structures, which was experimentally supported by negative-stain EM. This structural organization supports multiple, non-exclusive roles possible for MISHA. First, in line with recent proposals [96], MISHA may act as a chaperone or holdase, stabilizing protein or lipid components, via its hydrophobic groove [96]. Second, we propose the possibility of a previously unrecognized mechanism for forming cristae in which MISHA dimers serve as an inverted membrane scaffold supporting the initial membrane invagination (protrusions) for cristae growth (cartoon insert, Fig. 5D). Putative self-assembly into higher order structures may further stabilize and shape the cristae tubule. While this potential mechanism resembles that of banana-shaped BAR domain proteins assembling tip-to-tip, there is only one known mitochondrial representative – CIBAR1, a matrix protein identified during the development of our screen [88]. Despite limited data for CIBAR1, its abundance correlated uniquely with MISHA, which is prominently expressed in the brain where CIBAR1 is essential for development [118]. MISHA also correlated with the MIC60 network, consistent with its reported proximity to MIC19 [119]. Collectively, these findings reinforce an overarching role for MISHA in impacting membrane shape, possibly in early cristae formation (upstream or jointly with MICOS) or as a general support factor (for example in chaperoning cristae-organizing machineries or lipids that we did not test).

Besides confirming its membrane-binding, sculpting, or potential chaperoning activities, resolving MISHA’s experimental structure, and its regulation, will be essential to elucidate its molecular mechanism linked to mitochondrial ultrastructure. More generally, our discovery of this IMS protein as a critical effector of mitochondrial cristae shape, will surely help explain its role in pathology, positioning MISHA as a compelling focus for future research.

Beyond the hits and biological insights obtained by our screen, our work brings forward hMultiCLEM as a unique approach for systematic ultrastructure visualization. hMultiCLEM now enables the capacity to routinely screen with EM. This revolution extends to diverse applications, since hMultiCLEM enables genetic and drug screening, as well as time- and dose-response studies in lab cell lines, with potential for patient cells and disease models. By minimizing sample preparation noise and boosting throughput, this method advances the EM field into a new era of multiplexed, scalable & comparative, EM for membrane morphology studies in basic cell organelle research and medical contexts.

## Author Contributions

Sarah Hassdenteufel: Conceptualization; data curation; formal analysis; validation; investigation; visualization; methodology; writing – original draft. Yury Bykov: Methodology. Benjamin Dubreuil: Formal analysis; visualization. Katja Noll: Investigation. Ofrah Faust: Investigation. Roman Kamyshinsky: Investigation. Maxim Itkin: Investigation. Nir Cohen: Methodology. Sergey Malitsky: Investigation. Yoav Peleg: Investigation. Shira Albeck: Investigation. Karina von der Malsburg: Investigation. Deborah Fass: Conceptualization; resources; supervision; funding acquisition; project administration; Investigation. Martin van der Laan: Conceptualization; resources; supervision; funding acquisition; project administration. Maya Schuldiner: Conceptualization; resources; supervision; funding acquisition; project administration; writing – original draft. All authors contributed to the reviewing and editing of this manuscript.

In addition to the CRediT author contributions listed above, the contributions in detail are:

S.H. designed, performed, and analyzed the experiments and developed methodology. Y.B. designed and coded the hMC software workflow. B.D. curated and analyzed protein abundance and correlation data and analyzed lipidomics data. O.F. performed SEC-MALS analysis. K.v.d.M. designed, and K.N. performed the mitochondria enrichment with BN-PAGE analysis. M.I. and S.M. extracted and ran lipidomics samples and processed data. N.C. supported method development and provided training in CLEM. Y.P. and S.A. produced recombinant protein. D.F. produced recombinant protein, designed and performed the PEG-MAL assay, and designed and performed jointly with R.K. the negative-stain EM analysis. S.H. and M.S. wrote the manuscript, which all authors reviewed and provided feedback on. M.v.d.L. and D.F. provided scientific guidance and secured funding. M.S. supervised the work and secured funding.

## Acknowledgments and Funding Sources

We thank all lab members that read the draft, Ofir Klein and Rosario Valenti for their valuable expertise, Amir Fadel and Yeynit Asraf for expert technical support, Prof. Oliver Daumke and Prof. Jan Riemer for discussions on MISHA, and the EMBO EM Course 2022 for discussions on hMultiCLEM (and especially Prof. Yannick Schwab, Prof. Petr Chlanda, Prof. Gareth Griffiths, Prof. Andreas Brech, Jens Wohlmann, Urska Repnik). We are grateful to the Weizmann Institute of Science Electron Microscopy unit for their support (and especially to Nili Dezorella, Sharon Wolf and the late Eyal Shimoni) and to the Weizmann Institute of Science Stem Cell Core and Advanced Cell Technologies Unit, LCSF, for providing the Seahorse facility (and especially to Elena Ainbinder and Kira Orlovsky).

SH was supported by a postdoctoral scholarship from the Israel Academy of Sciences and Humanities (IASH). SM and MI work is supported by the Vera and John Schwartz Family Center for Metabolic Biology (Weizmann Institute of Science, Rehovot, Israel).

Work in the Schuldiner lab is supported by an SFB1190 grant from the Deutsche Forschungsgemeinschaft (DFG), the Irving and Cherna Moskowitz Center for Nano and Bio-Imaging (Weizmann Institute of Science), and a DFG Middle-Eastern collaboration grant (1028/11-1). The screening setup in the Schuldiner lab was purchased through the kind support of the Blythe Brenden-Mann foundation. MS is an Incumbent of the Dr. Gilbert Omenn and Martha Darling Professorial Chair in Molecular Genetics.

## Materials and Methods

### Cell culture and siRNA-mediated gene silencing

HeLa cells (CCL-2, ATCC) were cultivated at 37°C in DMEM GlutaMAX, 4.5 g/l glucose and 1 mM sodium pyruvate (10569010, Thermo Fisher Scientific), supplemented with 10% FBS (Thermo Fisher Scientific) and 1% penicillin-streptomycin (Biological Industries) in a humidified environment with a 5% CO_2_ atmosphere. Cells were frequently tested for mycoplasma contamination by PCR. Cells were passaged 24 hours prior to treatments. For pre-screening of growth conditions prior to screening of candidate proteins, HeLa cells were cultured in medium containing low (670086, Gibco EMEM; 1 g/L glucose, 1 mM sodium pyruvate, 2 mM L-glutamine) or high glucose concentrations (4.5 g/l), optionally supplemented with glutamine (11960044, Gibco DMEM without L-glutamine; 41965039, Gibco DMEM, 2 mM L-glutamine; or 10566016, Gibco DMEM GlutaMAX) and pyruvate (1 mM, 11360039, Gibco; or 10569010, Gibco DMEM GlutaMAX, 1 mM pyruvate). For 72 hours gene silencing prior to barcoding or Western Blot analysis, 2×10^5^ HeLa cells were seeded into a 60 mm culture dish with standard culture medium and treated with a high-complexity siRNA pool targeting a single gene or a non-targeting control (siPOOL synthesized by siTOOLs Biotech; see Supplementary table S1) to a final concentration of 3 nM using Lipofectamine RNAiMAX (13778075, Thermo Fisher Scientific) according to the manufacturer’s instructions. For 48 hours and 96 hours gene silencing, 4×10^5^ and 1.5×10^5^ HeLa cells were seeded into a 60 mm culture dish, respectively. For 72 hours gene silencing prior to Blue-native PAGE analysis, 1,95×10^6^ HeLa cells were seeded into a 150 mm or 100 mm culture dish. For 72 hour gene silencing prior to lipidomics analysis, 7×10^5^ HeLa cells were seeded into a 100 mm culture dish. For 72 hours gene silencing prior to Seahorse analysis, HeLa cells were seeded into Seahorse XF96 Cell Culture Microplates (103793-100, Agilent) at a density of 1,9×10^3^ cells per well. For 72 hours gene silencing prior to MitoTracker™ Orange staining, HeLa cells were seeded into 96-well glass-bottomed microscopy plates (Matrical Bioscience) at a density of 3×10^3^ cells per well. The amount of Lipofectamine RNAiMAX used for transfection was adjusted according to the number of cells seeded. For harvesting prior to barcoding, Western Blot or Blue-native PAGE analysis, cells were detached by treatment with low amounts of 0.25% trypsin (Bio-lab) for 2 minutes at 37°C and resuspended in culture medium.

### Cell surface barcoding

Prior to staining, harvested cells were washed three times with HBSS buffer (14175095, Thermo Fisher Scientific), gently centrifuged at 135 g for 2 minutes, and the supernatant was removed. For barcoding, WGA-conjugated CF™ dyes (29027-1, CF™ 405S; 29022-1, CF™ 488A; 29076-1, CF™ 555; 29026-1, CF™ 640R; Biotium) were dissolved in distilled water to prepare a 2 mg/ml stock solution (in PBS). Staining solutions were then prepared by diluting the dyes in HBSS buffer at concentrations determined by prior titration (see Supplementary Fig. 1). Cell pellets were resuspended in staining solution, ensuring that the cell concentration did not exceed 1 × 10⁶ cells per 100 µl, as recommended by the manufacturer and verified using a Countess cell counter (Invitrogen). After incubation at 37 °C for 10 minutes with gentle shaking, cells were gently pelleted and washed three times with HBSS buffer to remove residual dye. Stained cell populations were either immediately processed for EM sample preparation and kept separate until pooling for high-pressure freezing, or evaluated by fluorescence imaging in 96-well glass-bottom microscopy plates (Matrical Bioscience) using HBSS buffer.

### EM sample preparation

Barcoded HeLa cells were pooled to a single sample and gently sedimented to remove the supernatant. Preparation and embedding of barcoded HeLa cell pellets were performed using an in-resin CLEM protocol [47,49]. Briefly, a drop of viscous cell suspension was transferred into the 0.1 mm-deep cavity of a 0.1/0.2 mm membrane carrier (aluminum or gold, Martin Wohlwend) for high-pressure freezing. The cavity was then covered with the flat side of a 0.3 mm carrier, forming a specimen sandwich. The assembly was inserted into the Leica ICE high-pressure freezing machine using the standard cartridge system. Vitrified samples were subsequently stored under liquid nitrogen conditions. Resin embedding was carried out using a Leica AFS2 freeze-substitution system equipped with a processing robot. Samples were embedded in Lowicryl HM20 resin (14340, Electron Microscopy Sciences) following the freeze-substitution and embedding protocol optimized for in-resin CLEM [49]. Extra dry acetone (326801000, Thermo Fisher Scientific) containing 0.1% (w/v) uranyl acetate (UA), prepared from a 5% UA in methanol stock (EMS), was used as freeze-substitution medium. Resin blocks were trimmed using a Diatome 45° trimming knife, and 100 nm-thick sections were cut using a Diatome 35° ultra knife on a Leica UC7 ultramicrotome. Sections were mounted on 200-mesh copper grids with continuous carbon support film (CF200-Cu-TH, Electron Microscopy Sciences). Prior to EM imaging, grids were triple post-stained: 1 minute in Reynolds’ lead citrate (EMS), 10 minutes in a 2% UA aqueous solution (prepared from a 4% stock, EMS), and 1 minute in Reynolds’ lead citrate.

### Seahorse mitochondrial respiration analysis

Whole-cell mitochondrial respiration was assessed using the Seahorse XF96 Analyzer (Agilent Technologies) and the Seahorse XFe96/XF Pro FluxPak (103793-100, Agilent). Prior to the assay, HeLa cells were washed twice and incubated in freshly prepared XF assay Medium (103575-100, Agilent; Seahorse XF Base Medium) supplemented with 1 mM pyruvate, 2 mM L-glutamine, 10 mM glucose at 37°C in a non-CO₂ incubator for 1 h to equilibrate. Oxygen consumption rate (OCR) was measured using the XF Cell Mito Stress Test by sequential injection of mitochondrial modulators (Oligomycin, 1.5 uM, O4876, Sigma-Aldrich; FCCP, 0-5 uM, C2920, Sigma-Aldrich; Rotenone, 0.5 M, R8875, Sigma-Alrich; Antimycin A, 0.5uM, A8674, Sigma-Aldrich). ATP production was calculated from the obtained parameters according to the manufacturer’s instructions. Upon completion, cells were washed and stained using the CyQUANT™ Direct Cell Proliferation Assay (10ug/ml, C35011, Invitrogen) and measured on a plate reader. OCR values were normalized to cell numbers using WAVE software (Agilent). Relevant parameters were analysed using Excel and visualized using GraphPad Prism.

### MitoTracker™ Orange live-cell staining

Mitochondrial network labeling was performed using MitoTracker™ Orange CMTMRos (M7510, Invitrogen). Prior to staining, the media was removed and the cells were washed twice before adding pre-warmed medium containing the dye at a final concentration of 50 nM. Following incubation at 37 °C for 15 minutes in the dark, cells were washed with pre-warmed medium to remove excess dye and immediately imaged in medium containing HEPES (Sartorius) at a final concentration of 10 mM.

### Fluorescence microscopy of live and resin-embedded cells

All microscopy of fluorescently labelled HeLa cells (barcoding and MitoTracker™ staining) was performed using the Olympus Lumencor Spectra X in widefield mode. A 60x air objective (air, 0.9 numeral aperture), LED illumination (Olympus) and appropriate filter cubes were used for detection of the emitted fluorescence: MitoTracker™ Orange (Green, Triple); WGA-CF405S (Violet, DAPI); WGA-CF488 (Cyan, Quad); WGA-CF555 (Green, Triple); WGA-CF642 (Red, Cy5). Images of microscopy well plates from respective channels were automatically acquired using Olympus ScanR software. Images of resin-embedded sample sections from four channels were automatically acquired using the Olympus cellSens software. EM grids were sandwiched with a drop of distilled water between two glass coverslips and mounted onto the 96-well plate stage using a custom 3D-printed holder. For overview mapping, images were collected from transmission and far-red channels to locate the section. A focus map was generated for the complete section prior to collecting images.

### Transmission electron microscopy of multiplexed samples

EM of resin-embedded sample sections was performed using the FEI Tecnai G2 F20 TEM microscope operating at 120 kV, equipped with a TVIPS TemCam-XF416 retractable 16 megapixel CMOS camera. SerialEM software with navigator tool for automated data collection at multiple positions within the same section was used in a multi-step workflow [120,121]. Following overview mapping of the grid, the region of interest was defined to cover the entire section area. Images were acquired at medium magnification (nominal 1700×) with 4×4 binning (final resolution: 1024 × 1024 pixels), corresponding to a pixel size of ~4 nm at the specimen level. Micrographs were stitched to large-field montages using SerialEM and then processed using the established hMultiClaimer (hMC) software package. Using the generated SerialEM navigator file, feedback imaging of identified cells was prepared by realigning the grid registration in SerialEM. To capture complete individual cells, a 3 × 3 image montage of micrographs was collected at high magnification (nominal 6500×), using unbinned (1×1) mode, corresponding to a calibrated pixel size of approximately 1 nm/pixel.

### Negative-stain electron microscopy of recombinant protein

Carbon-coated copper grids (300 mesh) were glow-discharged for 120s at 0.5 mbar and 18 W with Evactron CombiClean System (XEI Scientific). Recombinant MISHA was diluted from a concentrated stock to 60 micromolar protein in 50 mM sodium phosphate buffer, pH 7.0, and 10 mM copper sulfate, and this solution was further diluted 1:10 before 3 µl was applied to the grid. After ~30 seconds, the excess solution was blotted, the grid surface was washed by contact with a 0.5-mL water drop, and the grid was stained by three applications of 10 µl each 2% uranyl acetate. TEM imaging was performed on Tecnai T12 (Thermo Fisher Scientific) microscope equipped with TVIPS TemCam XF416 camera (Tietz) and operated at 120 kV, as well as on Talos Arctica G2 TEM (Thermo Fisher Scientific) equipped with Falcon 4i direct detector (Thermo Fisher Scientific) and operated at 200 kV.

### Image processing and analysis

For LM visualization, the image contrast was linearly adjusted in Fiji/ImageJ and the images were converted from 16-bit to 8-bit. Cell segmentation and quantification of fluorescence intensity before EM sample preparation were performed using Olympus ScanR software. Segmentation was carried out on a virtual channel generated by merging the fluorescent channels. Dually stained cells were identified by applying gating based on fluorescence intensity thresholds. For LM processing, fluorescence images from four individual channels were stitched into large-field montages using Olympus cellSens software. Following conversion to TIFF in Fiji/ImageJ, the resulting montages were then applied to the established hMultiClaimer (hMC) software package. For EM visualization in Fiji/ImageJ, whole cell micrographs were batch-converted to TIFF using a MATLAB script enclosed in the hMC package (applied any image adjustments (e.g., scaling, bit depth)). Galleries of mitochondria were assembled using Fiji/ImageJ by extracting 500 × 500 pixel regions from the large-field montages. Typically, 100-250 mitochondria from at least 20-50 different cells were examined for each sample.

### HMultiClaimer software package

The software pipeline was based on our previous MutiCLEM package developed for yeast cell analysis [47]. The pipeline is build using MATLAB (MathWorks) image processing and statistics toolboxes, and general commands to organize the data. IMOD functions were used to process EM data [120,122]. For HMultiClaimer we took advantage of newly available generalist cell segmentation algorithm Cellpose [123]. Cellpose was used to detect cells in both LM and EM data independently. Then the cells detected by both modalities were matched based on their coordinates within each grid square. This allowed us to perform correlations locally, process larger sample areas, and reduce the requirements for correlation precision compared to the yeast pipeline. Other steps of the pipeline were performed essentially as described before [Bykov, Cohen…2019]. Briefly, the EM grid was imaged using LM and a stitched montage was obtained from the microscope software (see above). Then the medium magnification montage map of the sample was acquired using SerialEM (see above) and the navigator file was saved along with the montage. Then the EM montage was stitched using IMOD. Cell segmentation was performed using Cellpose separately on LM and EM montages. Based on cell outlines segmented in the LM image, peripheral fluorescence signal was measured for each cell and normalized as before [Bykov, Cohen 2019]. The barcodes were assigned using k-means clustering of normalized intensities. Then the MATLAB control point selection tool was used to correlate LM and EM montages and transform them to the same coordinate system. In the EM montage, grid bars grid squares were segmented based on intensity and the cells coordinates sorted by grid squares. Then the coordinates were matched to LM cell coordinates on per grid-square basis. The visual quality control was performed using a simple graphical interface that shows contrast-adjusted LM and EM data for a number of cells with the same automatically determined barcode. The operator reassigns incorrect barcodes if this is possible and identifies the cells that have a corrupted barcode or poor preservation in the EM sample (low quality cells). The low-quality cells are excluded from further processing. The coordinates of the high-quality cells are added to the SerialEM navigator file as new items using the built-in option for feedback imaging [ref Schorb]. Navigator item IDs contain the information about barcodes, square numbers, IDs of each cell. The code and the documentation for HMultiClaimer is freely available (https://mayaschuldiner.wixsite.com/schuldinerlab/lab-data).

### Protein extraction and Western blot analysis

SiRNA pools used for gene silencing were validated by western blotting. Upon 48-96 h silencing, cells were harvested, lysed and subjected to SDS-PAGE, western blotting and fluorescent-based imaging as described previously [12]. The following primary antibodies were used: rabbit anti-OPA1 (1:1000; 67589, D7C1A, Cell Signaling Technology), rabbit anti-ATP synthase subunit e (1:1000, 16483-1-AP, Proteintech), rabbit anti-CHCHD3/MIC19 (1:2000, HPA042935, Atlas Antibody), rabbit anti-APOOL/MIC27 (1:000, HPA000612, Atlas Antibody), rabbit anti-MIC60 (1:000, 10179-1-AP, Proteintech), rabbit anti-C1ORF151/MIC10 (1:500, ab84969, Abcam), rabbit anti-FAM136A (1:500, 30845-1-AP, Proteintech), and mouse anti-β-actin (1:2000; 8224, Abcam), served as a loading control. Fluorescently labeled secondary antibodies were: IRDye 800CW goat anti-rabbit IgG (1:10,000, 925-32211, Li-COR); IRDye 680RD goat anti-mouse IgG (1:15,000, 26-68070, Li-COR). Data were normalized to beta-actin and to control cells treated with non-targeting siRNA.

### Blue native PAGE analysis

Upon 72 hours gene silencing, harvested HeLa cells were washed twice with PBS (Sartorius) by gently centrifuging at 135 g for 2 minutes. The supernatant was removed, and the resulting pellets were shock-frozen in liquid nitrogen prior to mitochondria isolation. Frozen cell pellets were resuspended in buffer A (83 mM sucrose, 10 mM HEPES, pH 7.2) and homogenized with 10 strokes in a glass homogenizer. An equal volume of buffer B (250 mM sucrose, 30 mM HEPES, pH 7.2) was then added, followed by an additional 20 strokes of homogenization. The homogenate was centrifuged at 1,000 x g for 5 min at 4 °C, and the resulting supernatant was centrifuged at 12,000 x g for 5 min at 4 °C. The supernatant was discarded, and the pellet was resuspended in a small volume of buffer C (320 mM sucrose, 1 mM EDTA, 10 mM Tris-Cl, pH 7.4). Protein concentration was determined using the BCA Reagent ROTI®Quant. Aliquots of mitochondria were slowly frozen at −80 °C and stored until use.

Mitochondrial protein complexes were analyzed by blue native PAGE (BN-PAGE). Thawed mitochondria were solubilized on ice for 30 min in buffer containing 1% (w/v) digitonin, 20 mM Tris-HCl (pH 7.4), 0.1 mM EDTA, 50 mM NaCl, 10% (v/v) glycerol, and 1 mM PMSF. The lysate was centrifuged at 15,000 x g for 10 min at 4 °C to remove insoluble material. The supernatant was mixed with loading dye (5% Coomassie Brilliant Blue G in 500 mM ε-aminocaproic acid, 100 mM Bis-Tris, pH 7.0) and applied to a 4–13% BN-PAGE gel. Separated protein complexes were subsequently analyzed by Western blotting. The following primary antibodies were used: mouse anti-OPA1 (1:800; 612607, BD Biosciences), rabbit anti-MIC60 (1:500, raised in the van der Laan laboratory), rabbit anti-MIC10 (1:500, raised in the Pfanner/van der Laan laboratories), rabbit anti-ATP synthase subunit beta (1:500, raised in the Rehling laboratory). HRP-labeled secondary antibodies were either goat anti-mouse for OPA1 (Proteintech RGA001) or goat anti-rabbit (Proteintech RGAM001).

### Whole cell lipidomic analysis

Upon 72 hours gene silencing, HeLa cells were grown on a 100 mm cell culture dish and harvested in 0.5ml Methanol:DDW (1:1) through gentle scraping. Samples were immediately shock-frozen in liquid nitrogen and stored at −80°C until processing.

#### Lipids extraction

Extraction and analysis of lipids were conducted following previously established methods [81], with minor modifications. Approximately 2-5 million HeLa cells in 0.5ml Methanol:DDW (1:1) were lyophilized and extracted using 1 mL of a pre-chilled (−20°C) solution of methanol:methyl-tert-butyl ether (MTBE) at a 1:3 (v/v) ratio. Internal standards included phosphatidylcholine (17:0/17:0, Avanti; 0.1 µg/mL), phosphatidylethanolamine (17:0/17:0, Avanti; 0.4 µg/mL), Ceramide/Sphingoid Internal Standard Mixture II (Avanti, LM6005; 0.15 nmol/mL), d5-triacylglycerol Internal Standard Mixture I (Avanti, LM6000; 0.0267 µg/mL), and palmitic acid-13C (Sigma, 605573; 0.1 µg/mL). Samples were vortexed briefly, sonicated for 30 minutes in an ice-cold bath (with intermittent vortexing every 10 minutes), and then mixed with 0.5 mL of UPLC-grade water (DDW):methanol (3:1, v/v). After thorough vortexing and centrifugation, the upper organic phase was collected. The remaining polar phase was extracted using an additional 0.5 mL MTBE. Organic, extracts were combined and dried under a gentle nitrogen stream and stored at −80°C. Protein and debris were dried in a speedvac and was used for protein measurements for normalization.

#### LC-MS for lipidomics

Dried lipid extracts were resuspended in 120 µL acetonitrile:isopropanol (ACN:IPA, 75:25, v/v), vortexed, and centrifuged. Supernatants (100 µL) were analyzed using an Acquity I-class UPLC coupled with a Q Exactive Plus Orbitrap mass spectrometer. Chromatography employed an ACQUITY UPLC BEH C8 column (100 × 2.1 mm, 1.7 µm). Mobile phase A was DDW:ACN:IPA (46:38:16, v/v/v) containing 1% 1M NH4Ac and 0.1% acetic acid; phase B was DDW:ACN:IPA (1:69:30, v/v/v) with identical additives. The flow rate was 0.4 mL/min at 40°C. Gradient conditions: 100% A (0–1 min), decreasing to 25% A at 12 min, then to 0% A at 16 min, holding 100% B until 21 min, returning to 100% A at 21.5 min, with equilibration until 25 min. Injection volume was 4 µL.

Lipid data collection occurred in positive and negative switching HESI mode (m/z 100–1500) at capillary temperature 275°C, spray voltage 3.1 kV, sheath gas 60 units, auxiliary gas 20 units, and auxiliary gas temperature 300°C. MS1 resolution was 35,000 FWHM, MS/MS resolution 17,500 FWHM (1 m/z isolation).

#### Data analysis

Lipids were analyzed with LipidSearch™ software, validated against a custom lipid library, and the lipid elution order. Relative metabolite levels were normalized to internal standards and relative protein concentrations, assayed by Pierce™ BCA Protein Assay.

#### Quantification and Differential Analysis

Following whole cell lipidomics, mitochondrial lipid classes were quantified: mature cardiolipin (TLCL), monolysocardiolipin (MLCL), and phosphatidylglycerol (PG), and their constituent species across gene knockdowns. Normalized mass-spectrometry intensities were analyzed in R. For each knockdown, total class abundance was compared to its matched control using Welch’s *t*-test and Wilcoxon rank-sum test, with *p*-values corrected by the Benjamini–Hochberg method (FDR < 0.05). Log₂ fold-changes were calculated and visualized as grouped bar plots with confidence intervals and significance annotations (Fig. 4G).

To resolve species-level alterations, linear modeling with empirical Bayes moderation (LIMMA, [124]) was applied to log₂-transformed intensities. Differential abundances of mitochondrial lipid species upon gene knockdown were visualized as heatmaps (Supplementary Fig. 7B).

### Protein and mRNA Expression in HeLa Cells

Protein and mRNA abundance data for HeLa cells were respectively aggregated from the PaxDB integrated dataset [125] and the Human Protein Atlas (HPA) [126,127]. To enable cross-dataset comparison, all normalized abundance values were converted into expression percentiles. Expression percentiles for 70 human genes included in the hMultiCLEM screen were visualized as a heatmap to illustrate tissue-specific abundance patterns (Supplementary Fig. 2B).

### Protein Expression Analysis Across Human Tissues and Organs

Protein abundance data across a panel of 65 human tissues and organs were obtained from the integrated PaxDB resource [85]. We kept only 30 datasets where ≥10,000 proteins were detected covering at least 50 of the 70 hMultiCLEM candidates which include the following list of tissue and organs: Plasma, Heart, Liver, Esophagus, Prostate, Colon, Pancreas, Placenta, Skin, Salivary gland, Thyroid, Adrenal gland, Spleen, Cerebral cortex, Gallbladder, Rectum, Stomach, Lung, Kidney, Saliva, Testis, Lymph node, Bladder, Tonsil, Frontal cortex, Brain, Ovary, Uterus, Fallopian tube, and Platelet. Normalized protein abundances were converted to expression percentiles for each dataset similarly to cell line data. Tissues and organs were hierarchically clustered (Ward.D2 method) based on the pairwise Pearson correlation matrix of their complete proteome profiles, providing a biologically meaningful tissue order independent of the subset of proteins analyzed.

Heatmaps were generated to visualize tissue-based expression patterns of MICOS subunits, top screen hits, and mitochondrial proteins with proposed membrane-shaping activity on model membranes (Fig. 5A and Supplementary Fig. 8).

### Co-expression and Correlation Analysis

To assess potential functional relationships, co-expression analysis was performed using tissue expression percentile data for MICOS subunits, top screen hits, and mitochondrial proteins with proposed membrane-shaping activity (defined by liposome tubulation assays). Pearson correlation coefficients were calculated across the relative expression profiles of these 22 proteins, and the resulting correlation matrix was visualized as a correlogram, with proteins hierarchically clustered using complete linkage and the Ward.D2 method.

### Visualization and Assignment of Cell Barcodes with Uniform Manifold Approximation and Projection

Uniform Manifold Approximation and Projection (UMAP, [128]) was employed both to visualize high-dimensional fluorescence data from single cells and to predict experimental barcodes based on their fluorescence signatures. The workflow comprised three main stages: (i) visualization of single-cell fluorescence profiles in reduced dimensions, (ii) barcode assignment from UMAP embeddings, and (iii) quantitative evaluation and optimization of UMAP parameters to improve classification accuracy.

#### Visualization of Single-Cell Fluorescence Profiles

Normalized four-channel fluorescence intensity profiles from individual cells were projected into a two-dimensional space using UMAP. This dimensionality reduction from four to two dimensions enabled intuitive visualization of cell populations according to their fluorescence patterns to facilitate barcode assignment.

#### Predictive Barcode Assignment Using UMAP

Following dimensionality reduction, K-means clustering (*k* = 15) was applied to the UMAP coordinates to group cells according to the 15 known experimental barcodes. Idealized barcode vectors (e.g., [1, 0, 0, 0]) representing theoretical fluorescence profiles were transformed into the same UMAP space using the trained embedding model, producing reference positions for each barcode. Cluster centroids were compared with these projected barcode positions, and each cluster was assigned to the nearest theoretical barcode.

#### Evaluation and Optimization of UMAP Parameters

Since the quality of UMAP embedding influences both visualization clarity, cluster separation and subsequently barcode classification, a systematic grid search was performed across UMAP hyperparameters (number of neighbors = 15–200; minimum distance = 0.01–0.8; distance metric = Euclidean or Cosine). Each parameters configuration was evaluated using a complementary metrics, the average silhouette score (Equation 1).

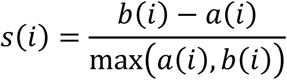

where *a(i)* is the mean distance from cell *i* to other cells in the same cluster, and *b(i)* is the mean distance to cells in the nearest neighboring cluster. Higher values indicate better cluster separation. The optimal parameters (*n_neighbors* = 200, *min_dist* = 0.01, *dist* = euclidean) yielded an average silhouette score of 0.78. These settings were used for the final UMAP embedding and barcode assignment shown in Fig. 2E.

### AlphaFold 3 predictions

Protein structure predictions were performed using AlphaFold 3, employing multiple protein copies [94]. The top-ranked model based on the predicted Template Modeling (pTM) score, was chosen for analysis. Structural confidence metrics, including pTM and interface pTM (ipTM), as well as piDDT (predicted inter-residue Distance Difference Test) were assessed to evaluate model reliability. All visualizations were carried out using ChimeraX 1.8 (UCSF).

### Quantification and statistical analysis

Protein levels from 3 to 6 repeats were quantified using Fiji/ImageJ. Statistical analysis using unpaired, two-tailed Student’s t-tests and visualization was performed in Excel. Total lipid class levels from 4 repeats were statistically analyzed using the Welch’s t-test, while for the analysis of individual lipid species, the limma package in R was used. P-values were adjusted for multiple testing using the Benjamini–Hochberg method. Lipidomics data were visualized in R. Respiration data from two runs with 3 to 6 repeats were analyzed using WAVE software (Agilent) and Excel, and visualized using GraphPad Prism. Samples from western blotting, lipidomics, and respiration analysis were compared to control cells treated with non-targeting siRNA Pool. Statistical significance is indicated in figures as follows: *p < 0.05; **p < 0.01; ***p < 0.001. Expression data were visualized in R. UMAP embedding and projection were carried out using the uwot package and visualized in R. K-means clustering and silhouette score calculation were performed with functions from the base stats and cluster packages, respectively. The Hungarian algorithm for optimal barcode assignment was implemented via the clue package. All figures in R were generated using the ggplot2 package and its extensions.

### Cloning, expression, and purification of FAM136A/MISHA

The human FAM136A/MISHA (Uniprot Q96C01, 138 amino acids) was expressed in the same bacterial host system from two tagged constructs: a SUMO-tagged version for biochemical and biophysical assays, and an MBP-tagged version for biochemical and biophysical analyses and negative-stain TEM.

#### SUMO-tagged protein

FAM136A/MISHA was cloned into the pET28-bdSumo vector using Restriction-Free (RF) cloning [129], yielding an expression construct encoding an N-terminal His₆–bdSUMO–FAM136A fusion. The construct was transformed into E. coli SHuffle® T7 Express cells (NEB, C3029J) and cultured in lysogeny broth (LB) supplemented with 30 µg/mL kanamycin. A 5 L culture was grown at 30 °C until reaching an optical density at 600 nm (OD₆₀₀) of 0.6–0.8, followed by induction with 200 µM isopropyl β-D-1-thiogalactopyranoside (IPTG). Protein expression was carried out for 16–18 h at 15 °C.

Cells were harvested by centrifugation and lysed using a cooled cell disruptor (Constant Systems) in lysis buffer containing PBS, 20 mM imidazole, lysozyme (200 KU/100 mL), DNase I (20 µg/mL), 1 mM MgCl₂, 1 mM PMSF, and a protease inhibitor cocktail. The clarified lysate was subjected to immobilized metal affinity chromatography (IMAC) on a HisTrap™ HP 5 mL column (Cytiva) using an ÄKTA FPLC system (Cytiva). The column was washed extensively with PBS containing 50 mM imidazole, and the His–SUMO–FAM136A fusion protein was eluted in PBS with 500 mM imidazole.

To remove the His-SUMO tag, His-bdSUMO protease (produced in-house) was added to the pooled fractions, and the mixture was dialyzed overnight at 4 °C against PBS containing 250 mM sucrose. The cleaved sample was re-applied to the same HisTrap column to separate the tag and protease from the cleaved FAM136A. The flow-through and 50 mM imidazole wash fractions contained the cleaved FAM136A (while the 500mM imidazole elution contained the tag and protease).

The cleaved FAM136A from the 50mM imidazole wash, was concentrated and further purified by size-exclusion chromatography (SEC) on a HiLoad™ 16/60 Superdex™ 75 column (Cytiva) equilibrated with 20 mM HEPES (pH 7.0), 150 mM NaCl. FAM136A eluted as a single peak (~55 mL). The pooled peak was diluted fivefold with 20 mM HEPES (pH 7.0) to remove the salt and applied to a HiTrap™ SP FF 5 mL cation exchange column (Cytiva) equilibrated with the dilution buffer. FAM136A was eluted using a linear NaCl gradient (0–1 M). Fractions containing pure FAM136A were pooled, concentrated to 1.5 mg/mL, aliquots were made which were flash-frozen in liquid nitrogen, and stored at −80 °C.

#### MBP-tagged protein

FAM136A/MISHA was produced as a fusion protein downstream of maltose binding protein (MBP) using a variant of the pMAL-C2 plasmid. The expressed protein contained a His6 tag, a TEV protease cleavage site, and an enterokinase cleavage site between the MBP and FAM136A. Cleavage with enterokinase was designed to yield FAM136A with the native methionine at the amino terminus. The plasmid was transformed into the Shuffle® E. coli strain (New England Biolabs). All incubations and culture growth occurred at 30 °C unless otherwise specified. Cultures (typically 1 L) were grown while shaking to 0.6 OD at 600 nm, at which point isopropyl β-D-1-thiogalactopyranoside was added to a final concentration of 0.5 mM to induce protein expression and the temperature was lowered to 24 °C. Cultures were harvested by centrifugation the following day, cell pellets were resuspended in phosphate buffered saline (PBS) containing 5 mM imidazole, and the suspensions were stored at −80 °C.

For protein purification, frozen cell suspensions were thawed at 4 °C and sonicated to produce homogenized lysate, which was then spun at 20,000 g. Supernatant was applied to a nickel nitrilotriacetic acid (Ni-NTA) column equilibrated in PBS plus 5 mM imidazole. After protein loading, the column was washed with the same buffer, and protein was eluted with a gradient of increasing imidazole. The eluted fusion protein was dialyzed against 25 mM Tris pH 7.5. After dialysis, 50 IU enterokinase (GenScript) was added to about 5 mL of about 200 µM fusion protein, and the cleavage reaction was allowed to proceed at room temperature for two days. Enterokinase cleavage was evaluated by SDS-PAGE, and if necessary to achieve at least 80% cleavage, another 20 IU enterokinase was added and incubation was extended for another day. The mixture was then re-applied to a Ni-NTA column equilibrated in PBS. The desired FAM136A product was eluted in batch mode at low imidazole concentrations (5 or 10 mM), whereas the MBP fragment containing the His6 tag eluted at higher imidazole (> 100 mM). FAM136A was concentrated using a centrifugal concentrator with 10 kD molecular weight cut-off and exchanged into 10 mM Tris, pH 7.5, 50 mM NaCl by eight dilution (at least 10-fold volume) and re-concentration steps. The protein concentration was estimated to be about 1 mM by diluting a sample 100-fold in the same buffer and measuring OD at 214 nm, using an extinction coefficient of 185,500 M-1cm-1 (https://bestsel-old.elte.hu/extcoeff.php).

### SEC-MALS

Recombinant MISHA/FAM136A was applied at a concentration of about 10 mg/ml to a Superdex 200 Increase 10/300 column (Cytiva) equilibrated in PBS, coupled to a DAWN HELEOS detector and an Optilab T-rEX differential refractive index detector (Wyatt Technology Corporation). The molecular weight was calculated using the ASTRA software package (Wyatt Technology Corporation). UV absorbance at 280 nm is missing due to limited aromatic amino acids in the MISHA sequence.

### Mass spectrometry

The LC/MS runs were performed on a Waters ACQUITY UPLC class H instrument, in positive ion mode using electrospray ionization. UPLC separation used a C4-BEH column (300 Å, 1.7 μm, 21 mm × 100 mm). The column was held at 40 °C and the autosampler at 10 °C. Mobile phase A was 0.1% formic acid in water, and mobile phase B was 0.1% formic acid in acetonitrile, and the flow rate was 0.4 mL/min. The MS data were collected on a Waters SQD2 detector with an m/z range of 600–1900 m/z. The desolvation temperature was 500 °C with a flow rate of 800 L/h. The voltages used were 1.00 kV for the capillary and 24 V for the cone. MassLynx version 4.2 was used to operate the LCMS and analyze the data. Raw data were processed using openLYNX and deconvoluted using MaxEnt with a range of 5000 : 30000 Da and a resolution of 1 Da/channel.

### Redox mobility shift assay

For reactions with maleimides, protein was diluted 25-fold into 10 mM Tris, pH 7.5, 50 mM NaCl. N-ethylmaleimide (NEM) (Sigma) was prepared at 1 M in acetonitrile, and MeO-PEG-mal (2 kDa) (Iris Biotech GmbH) (referred to here as PEG-mal) was dissolved at a concentration of 10 mM in ultrapure water. Both reagents were prepared fresh and used without delay. For the PEG-mal reaction under native conditions, 1 µL PEG-mal was added to 9 µL diluted protein and incubated for 10 minutes. Meanwhile, 1 µL NEM was added to 9 µL 4X SDS-PAGE loading buffer. At the end of the incubation with PEG-mal, 3 µL of the loading buffer containing NEM was added. For reactions under denaturing conditions, NEM was diluted 10-fold into ultrapure water, an aliquot was further diluted 10-fold into 4X SDS-PAGE loading buffer, and 3 µL were added to a 9 µL aliquot of diluted protein. Alternatively, PEG-mal was diluted 10-fold into SDS-PAGE loading buffer, and 3 µL were added to a 9 µL aliquot of diluted protein.

