## Supplementary Figures for "A multiplexed approach for genetic screening of human cells by electron microscopy uncovers a critical effector of mitochondrial cristae shape"

| (ug/ul) | B | G | R | FR |
|---|---|---|---|---|
| 1 | 90 | | | |
| 2 | | 30 | | |
| 3 | | | 30 | |
| 4 | | | | 60 |
| 5 | 180 | 30 | | |
| 6 | 180 | | 30 | |
| 7 | 180 | | | 60 |
| 8 | | 45 | 30 | |
| 9 | | 30 | | 60 |
| 10 | | | 30 | 60 |
| 11 | 180 | 30 | 30 | |
| 12 | 180 | 30 | | 60 |
| 13 | 180 | | 30 | 60 |
| 14 | | 45 | 30 | 60 |
| 15 | 180 | 30 | 30 | 60 |

**Supplementary Fig. S1: Determined dye concentrations for surface barcoding.**
Concentrations (ug/ul) of WGA-CF dyes (Biotium) used for combinatorial mixing and barcoding, determined by titration to achieve comparable fluorescence intensities for each label (1-15; see Fig. 1 and Materials and Methods). Four dyes/channels were used: WGA-CF405S/Blue (B), WGA-CF488A/Green (G), WGA-CF555/Red (R) and WGA-CF640R/Far-red (FR).
See Fig. 1C-D.

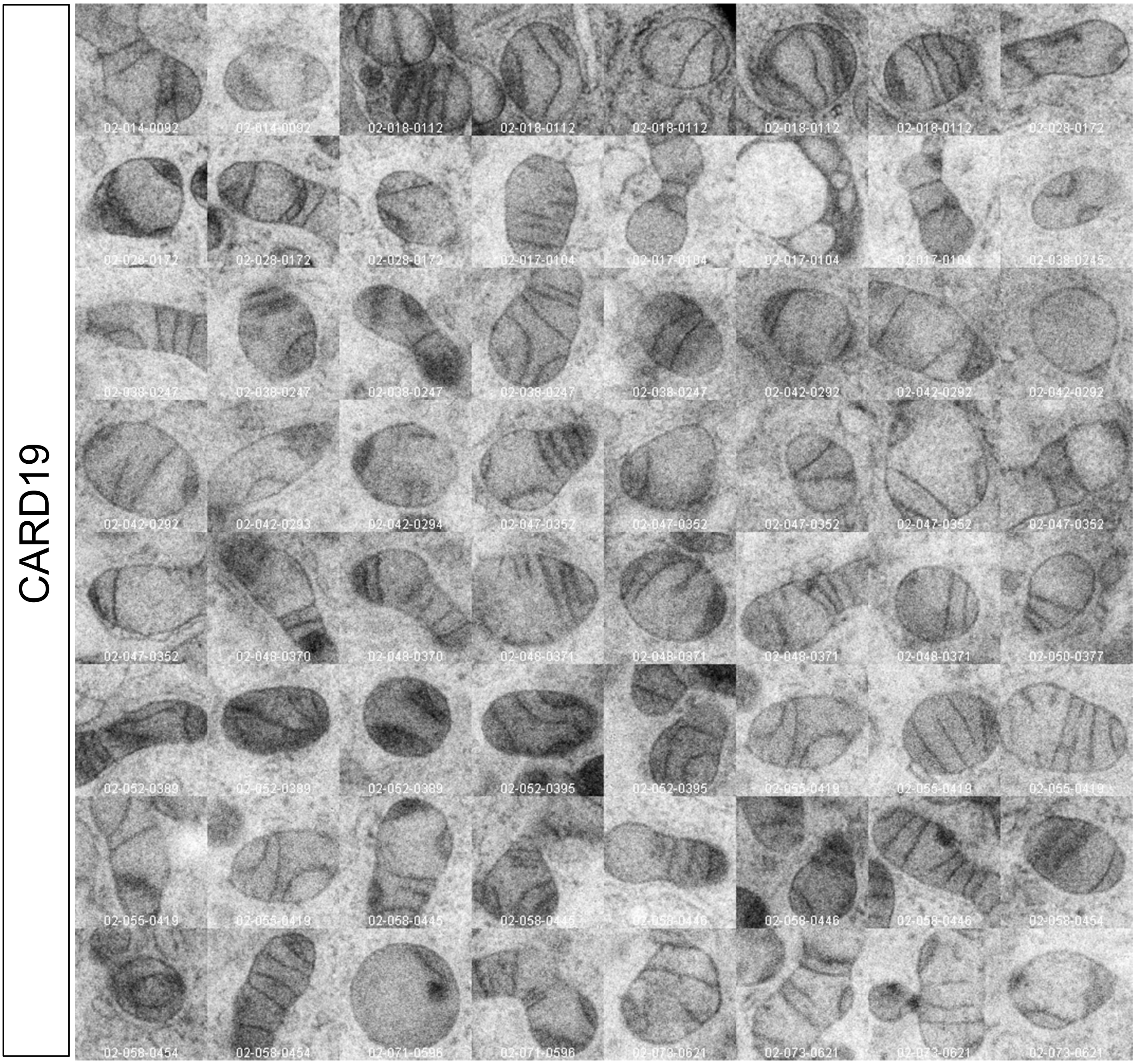


**Supplementary Fig. S2: Example gallery of ultrastructural features visualized by hMultiCLEM.**
Example of a low-resolution image gallery assembled in ImageJ to facilitate rapid visual comparison by browsing. Micrographs depict the phenotypic spectrum of mitochondria in cells subjected to 72 hours silencing and hMultiCLEM. See Figs. 2B and 4A.

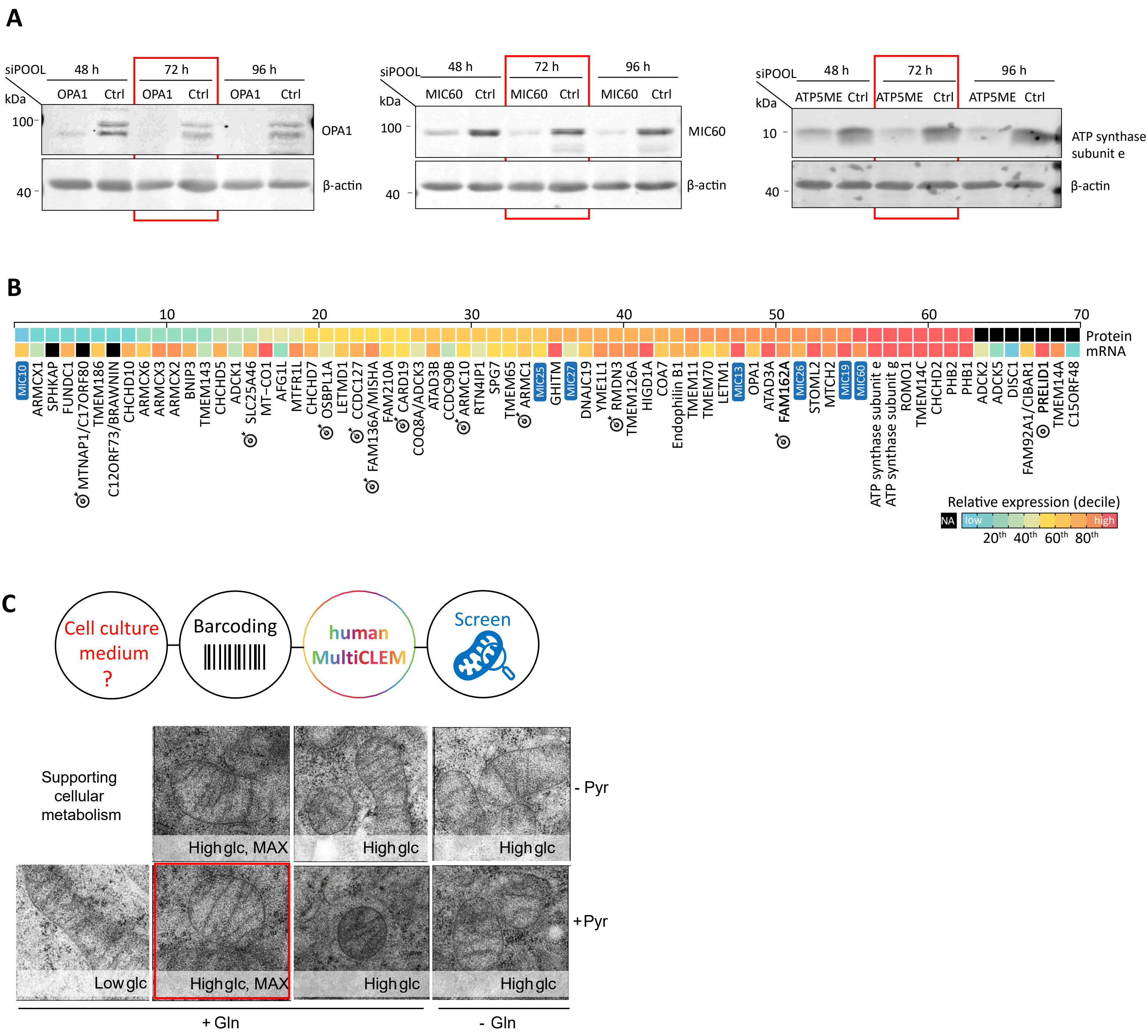


**Supplementary Fig. S3: Selected conditions for genetic screening.**

A) Depletion kinetics of OPA1, MIC60, and ATP5ME assessed by Western blot after 48, 72, and 96 hours of silencing. Red rectangle: selected condition for hMultiCLEM screen of candidate proteins.

B) Abundance of candidate proteins in Hela cells. A heatmap was generated for the 70 human genes selected for screening by hMultiCLEM, with expression percentiles color-coded from blue (lowest, 0–10th) to red (highest, 90–100th). Missing values are shown in black. MICOS subunits are annotated with a blue tag. Top candidates are highlighted (target symbol).

C) Pre-screening of culture media by hMultiCLEM. HeLa cells were cultured in media containing low or high glucose (Glc) concentrations, optionally supplemented with glutamine (Gln or DMEM GlutaMAX, indicated as MAX) and pyruvate (Pyr), as specified. Red rectangle: selected condition for hMultiCLEM screen of candidate proteins.

See Fig. 3B-D.

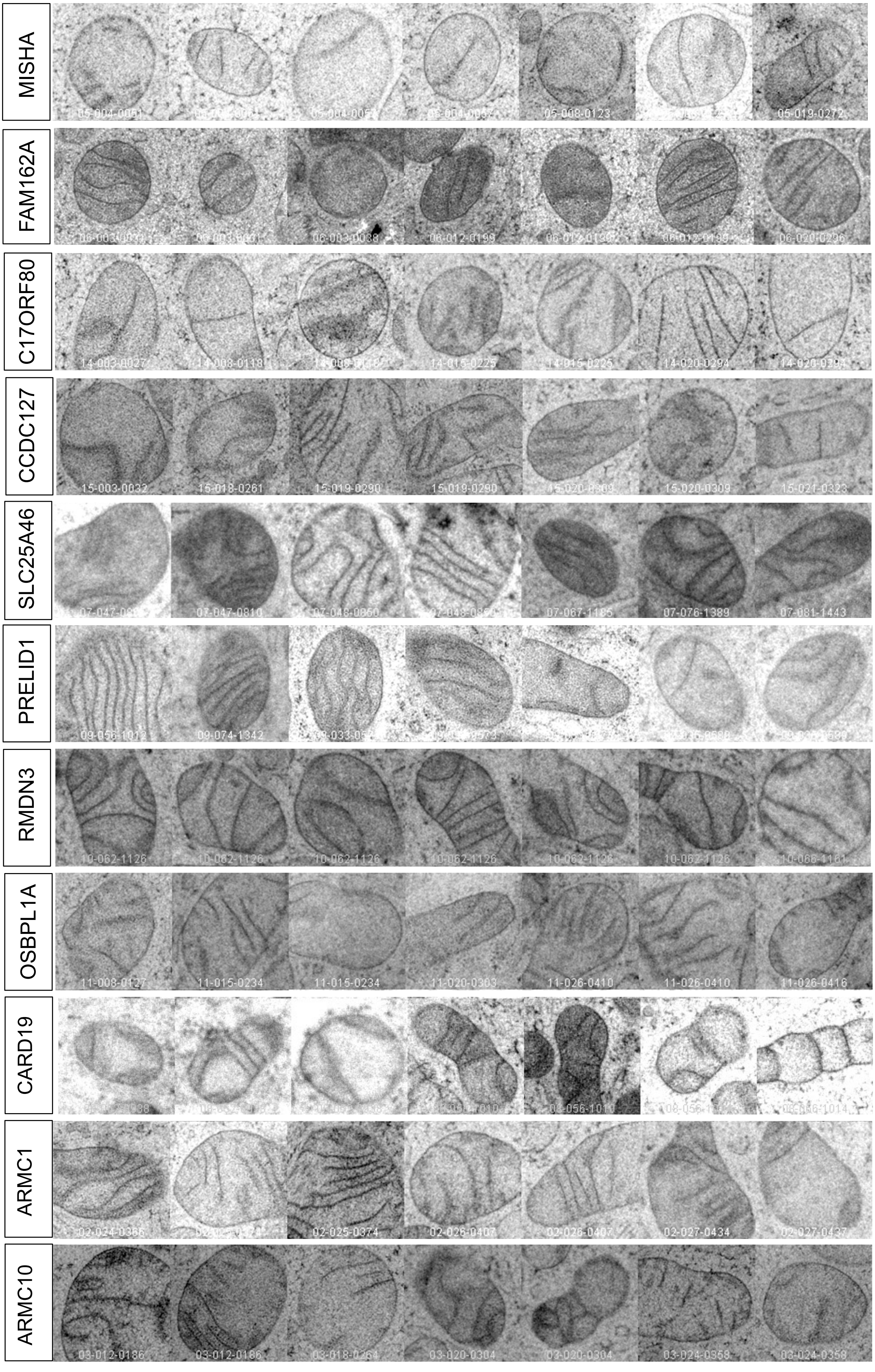


**Supplementary Fig. S4: Repeat and re-imaging of top candidates by hMultiCLEM.**
Small-scale repeat of the hMultiCLEM screen for top candidates after 72 hours of silencing. At least 50 cells per candidate were imaged. Micrographs depict representative mitochondrial phenotypes from randomly selected cells. Example of a low-resolution image gallery assembled in ImageJ to facilitate rapid visual comparison by browsing. See Fig. 4A.

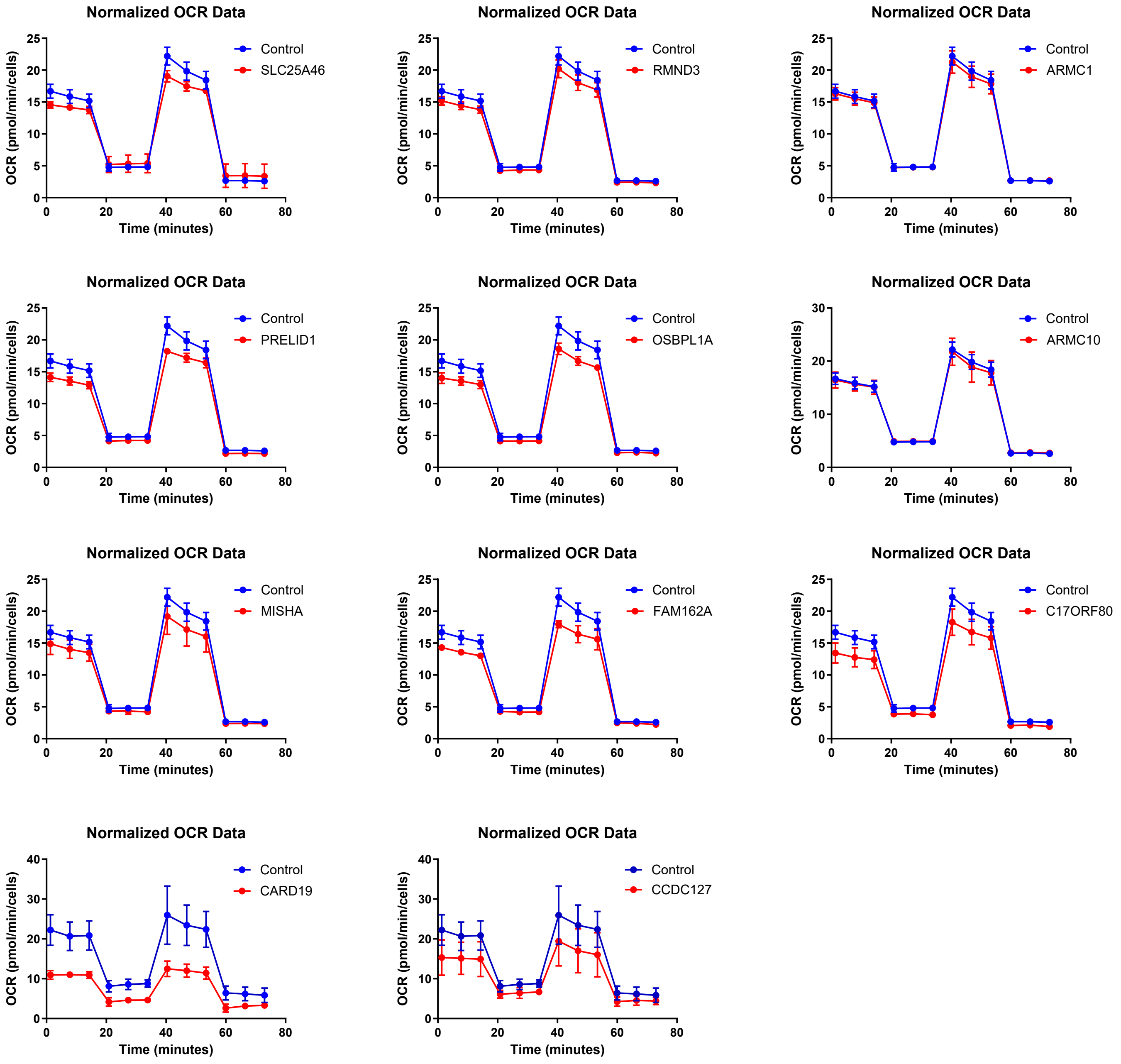


**Supplementary Fig. S5: Oxygen consumption rate profiles by Seahorse respiration assays.**
Upon 72 hours of silencing, HeLa cells were subjected to Seahorse respiration assay. The oxygen consumption rate (OCR) was measured at indicated time points along with application of OXPHOS inhibitors (see Fig. 4C). For parameters calculated from the measurements of 3-6 repeats, see Fig. 4D-E and Materials and Methods.

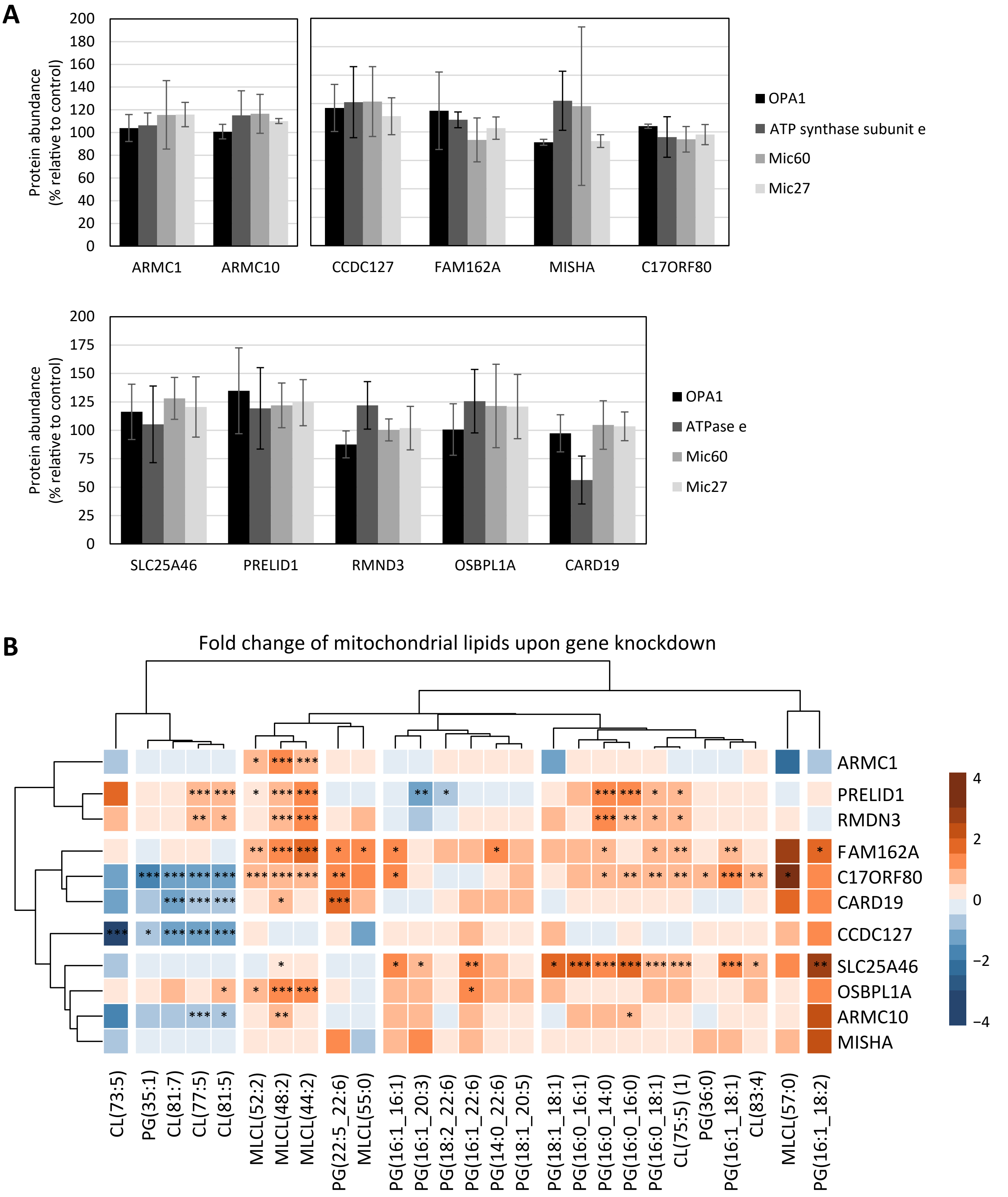


**Supplementary Fig. S6: Abundance analysis of core-cristae modulating proteins and individual lipid species.**
A) Upon 72 hours of silencing, HeLa cells were subjected to Western blotting. Bar graphs show mean ± SD, with statistical significance assessed by Student's t-test (n=3-6 replicates). For representative gel images, see Fig. 4F.

B) Upon 72 hours of silencing, HeLa cells were subjected to lipidomic analysis on whole cells. Data were normalized to the total protein content. Analysis of mitochondrial lipid species for mature cardiolipin (CL), monolysocardiolipin (MLCL) or phosphatidylglycerol (PG) are shown. The heatmap visualizes the $\log_2$ fold change ($Log_2FC$, calculated from means of 4 repeats) of indicated lipid species relative to control cells. Data were analyzed using linear modeling with empirical Bayes moderation (LIMMA). For analysis of total lipid classes, see Fig. 4G.
*$p<0.05$; **$p<0.01$; ***$p<0.001$.

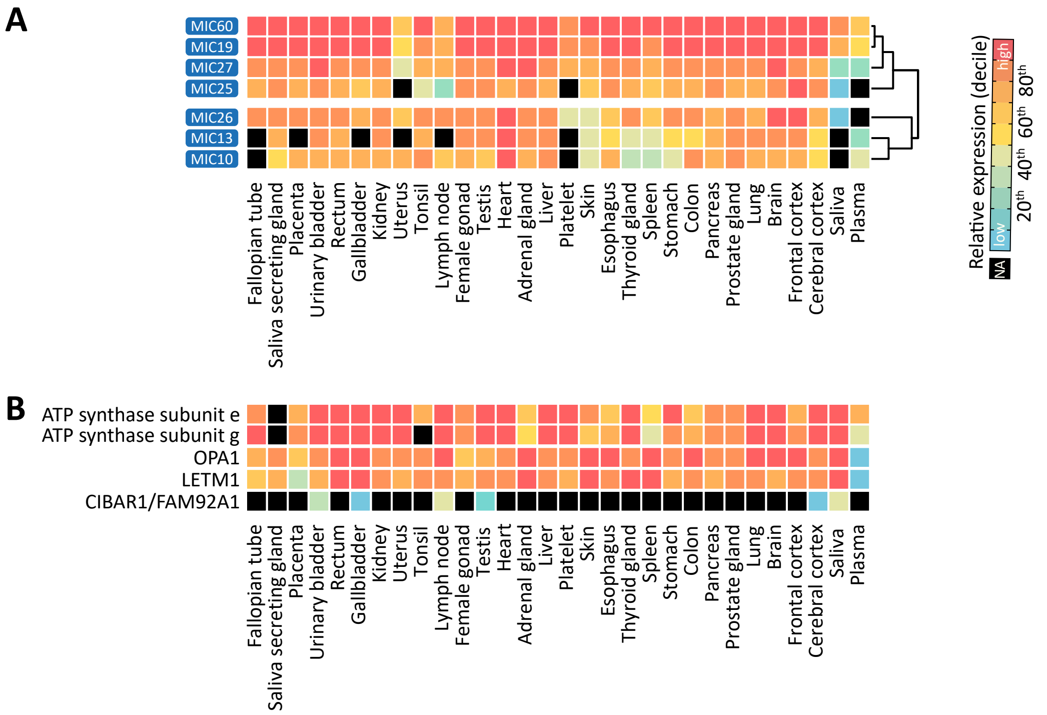


**Supplementary Fig. S7: Expression profile of MICOS components and membrane-shaping proteins across human tissues.**
A) Protein abundances of MICOS components. Black/NA: data not available in datasets. See Fig. 5 and Materials and Methods.

B) Protein abundances of ATP synthase dimerization subunits e and g, OPA1, LETM1 and CIBAR1. Black/NA: data not available in datasets. See Fig. 5 and Materials and Methods.

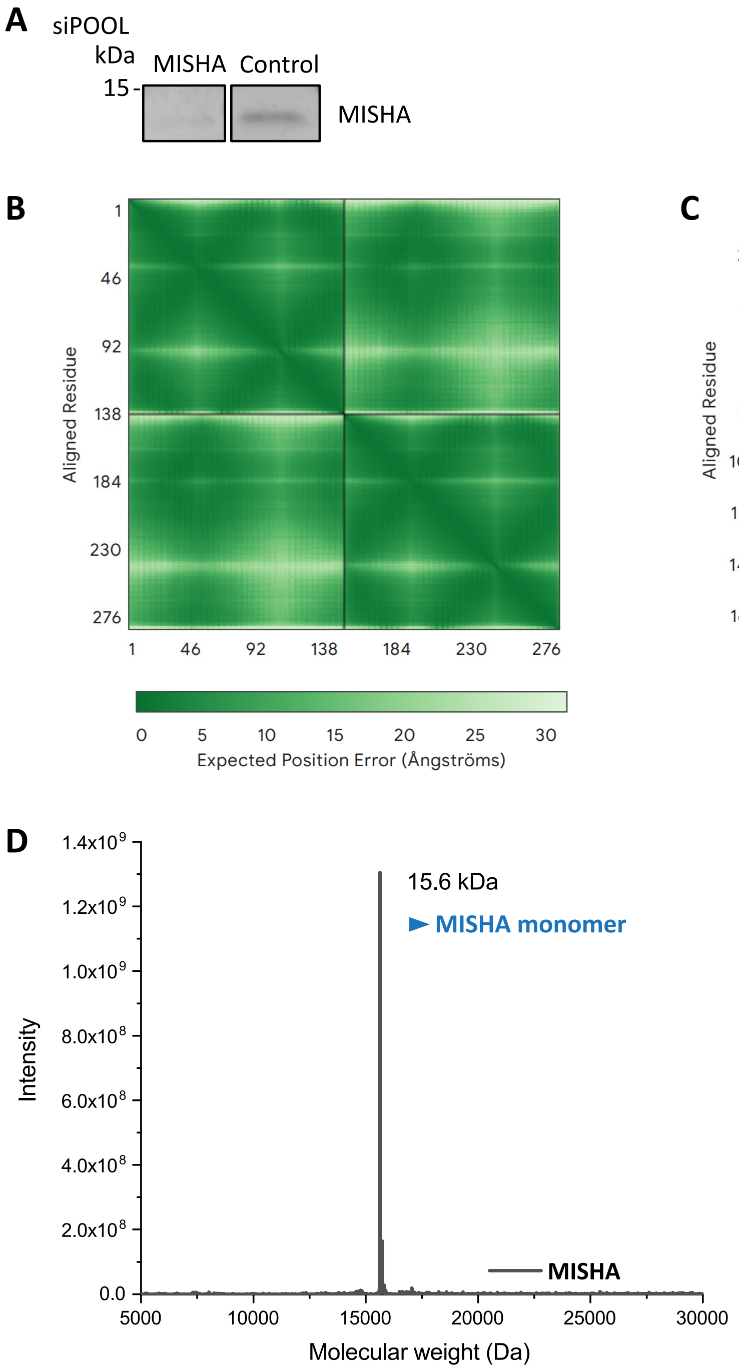


**Supplementary Fig. S8: The IMS protein MISHA forms a non-covalent dimer and is predicted to form higher-order assemblies.**

A) Depletion efficiency of MISHA in HeLa cells following 72 hours of silencing, as assessed by immunoblotting using an antibody against MISHA.

B-C) Confidence of AlphaFold 3 predictions for MISHA assemblies. The heatmaps display pLDDT scores (predicted Local Distance Difference Test), indicating the model's per-residue confidence (green = low, white = high). Enlarged views of Figure 6B and 6D.

B) Dimer.

C) Multimer of 12 protein copies.

D) Mass spectrometry analysis of recombinant MISHA.

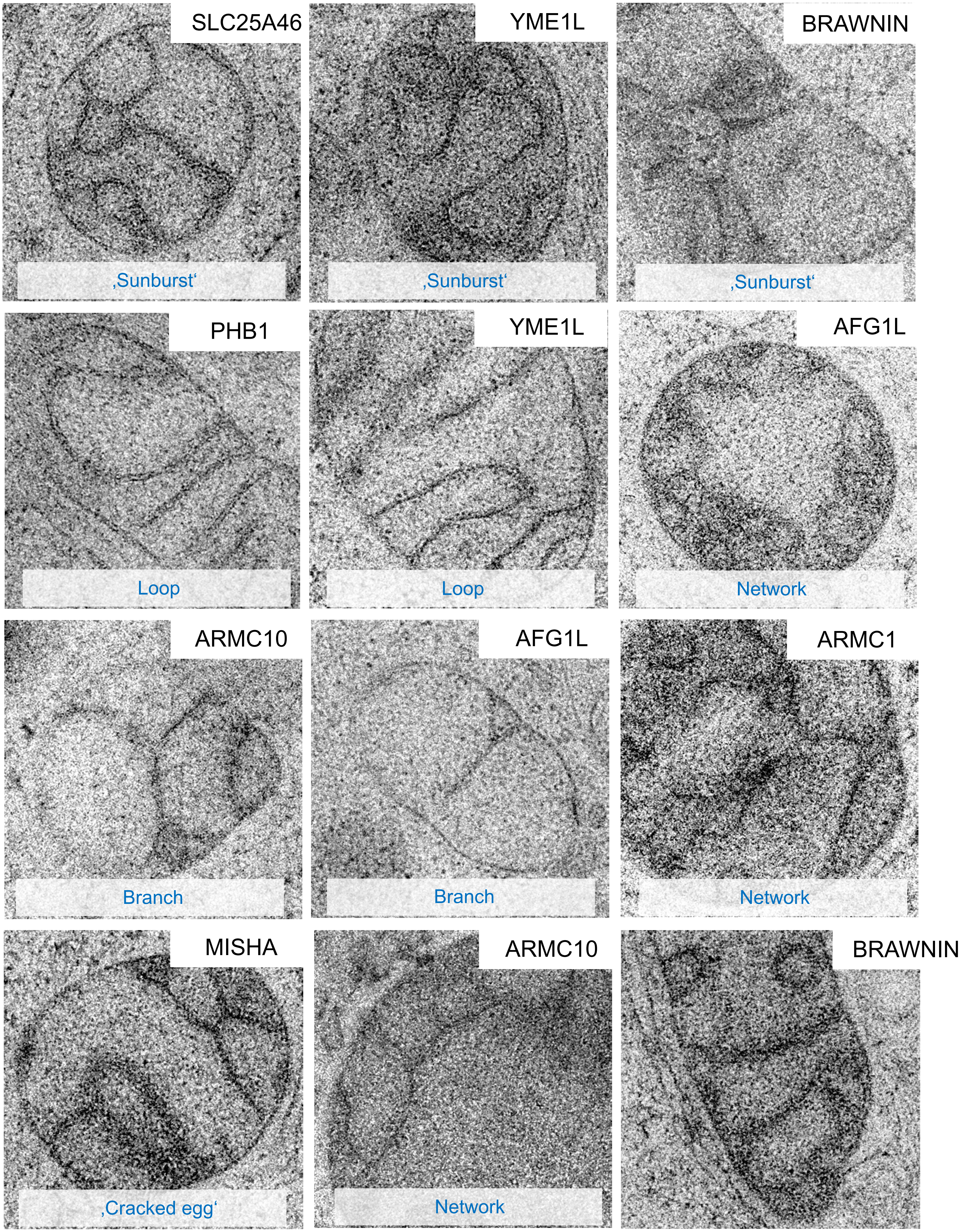


**Supplementary Fig. S9: Examples of rare cristae phenotypes by hMultiCLEM.**
Micrographs depict mitochondria of rare appearance observed in the hMultiCLEM screen following 72 hours of candidate silencing and absent in control cells. For example, cristae branches, networks (branched, swirled or radial (e.g. 'sunburst') and loops. Selected high-resolution images of Fig. 3E-F.

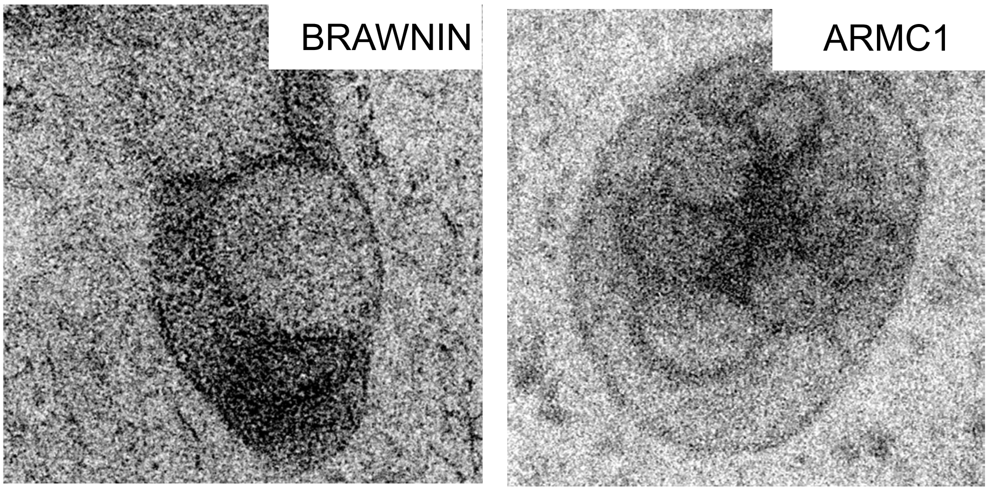
BRAWNIN
ARMC1
